# Activity-dependent modulation of NMDA receptors by endogenous zinc shapes dendritic function in cortical neurons

**DOI:** 10.1101/2021.09.17.460586

**Authors:** Annunziato Morabito, Yann Zerlaut, Benjamin Serraz, Romain Sala, Pierre Paoletti, Nelson Rebola

## Abstract

Activation of NMDA receptors (NMDARs) has been proposed to be a key component of single neuron computations in vivo. However is unknown if specific mechanisms control the function of such receptors and modulate input-output transformations performed by cortical neurons under in vivo-like conditions. Here we found that in layer 2/3 pyramidal neurons (L2/3 PNs), repeated synaptic stimulation results in an activity-dependent decrease in NMDARs activity by vesicular zinc. Such a mechanism shifted the threshold for dendritic non-linearities and strongly reduced LTP induction. Modulation of NMDARs was cell- and pathway-specific, being present selectively in L2/3-L2/3 connections but absent in ascending bottom-up inputs originating from L4 neurons. Numerical simulations highlighted that activity-dependent modulation of NMDARs has an important influence in dendritic computations endowing L2/3 PN dendrites with the ability to sustain dendritic non-linear integrations constant across different regimes of synaptic activity like those found in vivo. The present results therefore provide a new perspective on the action of vesicular zinc in cortical circuits by highlighting the role of this endogenous ion in normalizing dendritic integration of PNs during a constantly changing synaptic input pattern.

## Introduction

A fundamental computation performed by individual neurons in the brain involves the transformation of incoming synaptic information into an output firing pattern(Brunel et al., 2014; Silver, 2010). Synaptic inputs arrive primarily on the dendritic tree of neurons where a diversity of postsynaptic dendritic mechanisms in known to shape input-output relationships. These include synaptic saturation (Abrahamsson et al., 2012; Vervaeke et al., 2012), dendritic spikes(Häusser et al., 2000; Smith et al., 2013) as well as NMDAR nonlinearities(Major et al., 2013). However, how such dendritic mechanisms are actually recruited in vivo is much less understood.

In sensory cortices of awake mice spontaneous firing of single neurons can vary between low firing (<0.1 Hz) to considerably higher firing regimes (~10 Hz) depending on the behavioral and physiological state of the animal (McGinley et al., 2015; Vinck et al., 2015; Zerlaut et al., 2019). Theoretical studies suggest that such periods of increased spontaneous synaptic activity might facilitate recruitment of NMDARs and thus result in supralinear dendritic integration (Farinella et al., 2014; Ujfalussy et al., 2018). The increased activation of NMDARs is explained by their slow kinetics (half-decay 40-70 ms) that effectively summate over relatively long times integrating recent history of synaptic activity. However, in vivo activity patterns are likely to engage modulatory mechanisms (e.g. synaptic plasticity, firing adaptation, release of neuromodulators). Whether such modulatory mechanisms specifically apply to NMDAR function altering dendritic operations in cortical neurons is currently unknown. NMDARs are known to be particularly susceptible to regulation (Raman et al., 1996; Tong et al., 1995) and an eventual activity-dependent plasticity of synaptic NMDARs is expected to considerably impact integrative properties of single-neurons working under *in vivo*-like conditions.

Zinc is a metal ion that in the brain has the particularity of being co-stored together with glutamate inside synaptic vesicles of glutamatergic terminals through the action of a dedicated vesicular transporter (ZnT-3; Palmiter et al., 1996). During synaptic activity vesicular zinc is released into the extracellular space and has been shown to efficiently modulate synaptic NMDAR (Anderson et al., 2015, 2017; Assaf and Chung, 1984; Erreger and Traynelis, 2005; Howell et al., 1984; Pan et al., 2011; Paoletti et al., 2009; Sensi et al., 2009; Vergnano et al., 2014; Vogt et al., 2000). Such a modulatory system seems thus particularly well positioned to control NMDARs function during periods of increased network dynamics. Despite clear behavior alterations associated with manipulation of zinc signaling(Adlard et al., 2010; Anderson et al., 2017; McAllister and Dyck, 2017; Patrick Wu and Dyck, 2018) the mechanisms and computational role of vesicular zinc action in cortical circuits is understudied.

Here we combined two-photon (2P) imaging, slice electrophysiology, glutamate uncaging and numerical simulations to reveal that: (1) Zinc release modulates NMDAR in an activity-dependent and pathway-specific manner controlling dendritic integration in PNs in the primary somatosensory cortex (S1); (2) Zinc inhibition of NMDARs renders the recruitment of dendritic non-linearities particularly insensitive to previous history of synaptic activation. These results provide a new perspective for the role of zinc-containing neurons in brain microcircuits and reveal a mechanism by which neurons are able to normalize dendritic integration across different activity regimes often associated with the cortical states of wakefulness.

## Results

### Synaptic zinc is an endogenous neuromodulator of NMDAR function in L2/3 PNs

We hypothesized that a possible activity-dependent modulation of NMDARs might have important consequences in computational properties of cortical neurons. To test this, we focused our study in L2/3 PNs, known to express important NMDAR-dependent dendritic computations(Branco and Häusser, 2011; Major et al., 2013; Schiller et al., 2000) and to be particularly enriched in vesicular zinc, an endogenous negative modulator of NMDAR function (Figure 1A). We started by testing if synaptic NMDA-EPSCs were modulated during repeated synaptic activation. Interestingly, bath application of the zinc chelators tricine (10 mM; Paoletti et al., 1997) or ZX1 (100 μM; Anderson et al., 2015) revealed a use-dependent inhibition of NMDARs during trains of synaptic stimulation without noticeable effect in the amplitude of the first NMDA-EPSC (ratio amplitude 5th/1st pulse; control: 1.09 ± 0.05, Tricine:1.30 ± 0.05, n = 7, p = 0.008; control:1.10 ± 0.03; ZX1:1.30 ± 0.06, n = 10; p = 0.009, Wilcoxon matched-pairs signed rank test; Figure 1B,C and S1A). Importantly, at the concentrations used, both chelators can rapidly interfere with fast extracellular zinc elevations (Anderson et al., 2015; Paoletti et al., 2009; Vergnano et al., 2014) without affecting intracellular zinc levels (Figure S1B, C). Zinc modulation was also present at lower frequencies of stimulation (3 Hz) in agreement with slow dissociation from bound NMDARs (Figure S1D; Paoletti et al., 1997). The action of zinc chelators was absent in slices from ZnT3-KO mice that lack vesicular zinc (Figure S1E; Palmiter et al., 1996). Moreover zinc chelators had no effect in AMPAR-EPSCs, indicating that, as observed in hippocampal synapses (Vergnano et al., 2014) zinc effect is selective to NMDARs (Figure S1F,G). These results suggest that during trains of synaptic activity vesicular zinc specifically downregulates NMDAR function in L2/3 PNs.

**Figure 1:**
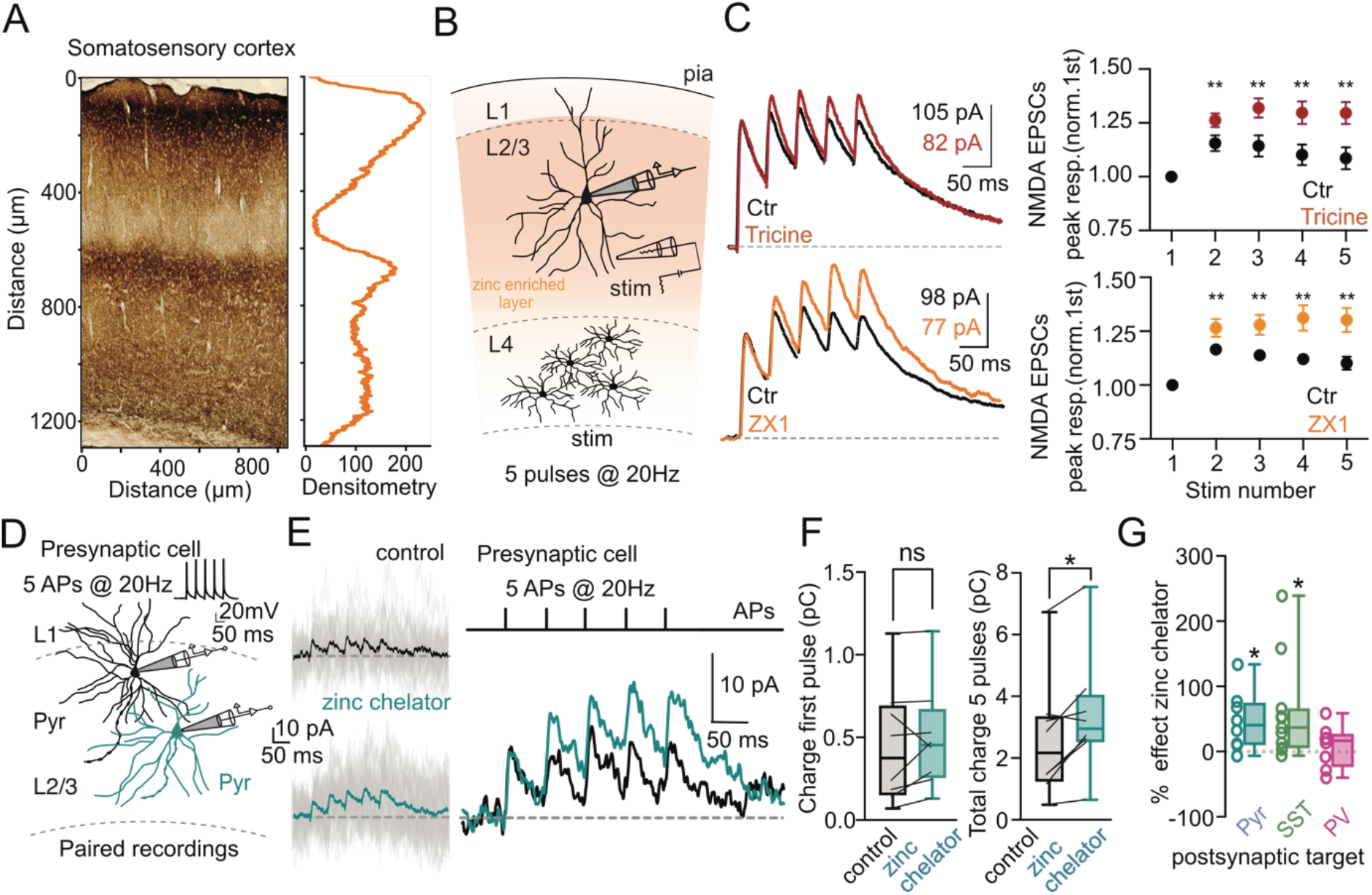
Zinc is an endogenous modulator of NMDAR function in S1. **A)** Timm’s staining (*left*) and corresponding densitometric profile (*right*). **B)** Schematic representation of experimental conditions used to stimulate local excitatory connections and record NMDA-EPSCs in L2/3 PN. **C**) Representative traces and summary plot of normalized NMDA-EPSCs peak amplitude during a train of synaptic stimulation (5 pulses @ 20 Hz) obtained in control conditions (black) and after bath application of zinc chelators tricine (red) and ZX1 (orange) in L2/3 PNs. Values are presented as mean ± SEM, * < p0.05 paired Wilcoxon matched-pairs test. **D)** Illustration of recording configuration used to record unitary NMDA-EPSCs between L2/3 PNs. Presynaptic cell was stimulated with a train of 5 APs (20 Hz). **E)** *Left*: Representative traces of uNMDA-EPSCs recorded in a postsynaptic L2/3 PN held at +30 mV. Single repetitions (30 sweeps) are in faint grey, average is in full color. *Right*: superimposed average uNMDA-EPSC obtained in control (black) and in presence of zinc chelator (dark cyan). **F)** Summary plot of effect of chelating extracellular zinc in the charge transfer carried by a train (5 APs @ 20 Hz) of uNMDA-EPSCs. **\***p<0.05, Wilcoxon matched-pairs signed rank test. Boxes represent interquartile ranges with the horizontal bars showing the medians. **G)** Summary plot of the effect of zinc chelators in charge transfer carried by a train of NMDAR-EPSCs at synapses between L2/3 PN and different postsynaptic targets (Pyr median: 40.45 %; SST: 36.76%; PV: 16.47%. Pyr: p = 0.026, SST p = 0.013, PV p = 0.90, Kruskal Wallis test compared to no effect).

L2/3 PNs receive inputs from local L2/3 and L4 neurons as well as from long range connections from other cortical regions. In order to measure the impact of zinc modulation at single synapses, we performed dual whole-cell recordings from synaptically connected L2/3 PNs (Figure 1D). In 8 tested pairs, chelating extracellular zinc significantly increased the total charge transfer carried by the train of NMDA-EPSCs by almost 50% (control = 2.61 ± 0.7 pC; chelator = 3.38 ± 0.7 pC; p= 0.023 Wilcoxon matched-pairs signed-rank test; Figure 1D-F). This effect was substantially larger than the effect obtained with extracellular stimulation compatible with the expected heterogeneous recruitment of zinc-positive and zinc-negative fibers with electrical stimulation. Again, zinc chelators did not alter charge transfer measured only during the first NMDA-EPSC in the train (control: 0.45 ± 0.13 pC; chelator: 0.48 ± 0.11 pC; p= 0.15, Wilcoxon matched-pair signed rank test; Figure 1D-F), arguing for a phasic and not tonic action of synaptic zinc in L2/3 PN NMDARs. Altogether these results indicate that zinc is an effective modulator of NMDARs at L2/3-L2/3 PN synapses.

### Cell- and pathway-specific actions of vesicular zinc in S1

The heterogenous labelling of free zinc in the cortex (Figure 1A) raised the question about the potential variability of the observed activity-dependent modulation of NMDARs across synaptic inputs. To evaluate such a possibility, we placed the stimulation electrode in L4, a cortical layer almost devoid of zinc staining, while recording NMDAR-EPSCs in L2/3 PNs (Figure S2A). Short-term depression was particularly evident when compared to local L2/3 stimulation (Figure 1C). This observation is compatible with the high-release probability previously reported at L4-L2/3 synapses (Feldmeyer et al., 2002; Silver et al., 2003). Interestingly, under these experimental conditions, zinc chelators did not modify amplitude nor the short-term plasticity of recorded NMDAR-EPSCs (Figure S2A). This effect was not due to a different sensitivity of postsynaptic NMDARs across the two inputs as exogenous zinc application (300 nM) produced similar degree of inhibition while evoking NMDA-EPSCs in L2/3 or L4 (Figure S2B). These results indicate that activity-dependent modulation of NMDARs across synaptic inputs is heterogeneous, potentially altering their contribution to synaptic integration and plasticity.

In addition to the variability between presynaptic inputs, we further investigated if modulation of NMDARs could also differ between postsynaptic targets. L2/3 PNs axons contact L2/3 PNs as well as other local interneurons (INs). For that we tested the impact of zinc chelators in uNMDARs-EPSCs recorded in somatostatin-positive (SST^+^)- or parvalbumin-positive (PV^+^)-INs. Extracellular zinc chelation significantly increased charge transfer carried by NMDARs in synapses between L2/3 PNs and SST^+^-INs (control: = 1.89 ± 0.55 pC; chelator: 2.73 ± 0.65 pC, n=11; p= 0.002, Wilcoxon matched-pair signed rank test; Figure 2G, S2C-F). Surprisingly, the same was not observed for synapses into PV-INs (Figure 2G, S2G-H). Unexpectedly in PV-INs, the lack of effect was associated with a reduced sensitivity of postsynaptic NMDARs to exogenous zinc application (Figure S2I-J). The underlying cellular mechanism behind such a difference in zinc sensitivity is at present unclear but is likely associated with variation in the subunit composition of synaptic NMDARs that can affect the sensitivity of synaptic NMDARs to zinc inhibition (Garst-Orozco et al., 2020; Paoletti, 2011).

**Figure 2:**
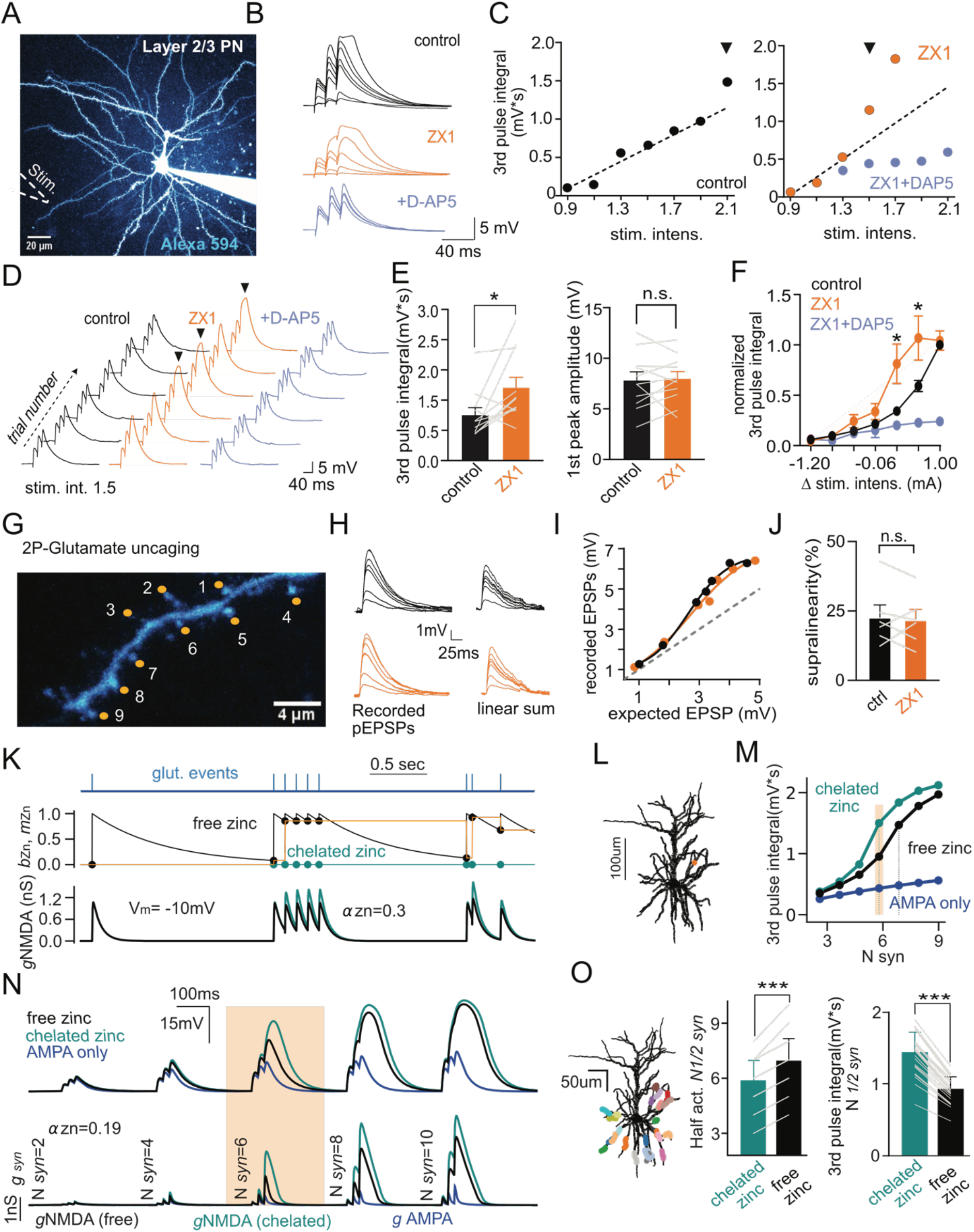
Synaptic zinc release impacts dendritic non-linearities in basal dendrites of L2/3 PNs. **A)** Two-photon laser scanning microscopy (2PLSM) image (maximum-intensity projection, MIP) of L2/3 PN. In white dotted line the location of theta glass pipette used for focal dendritic stimulation. **B)** Representative traces of recorded EPSPs for increasing (*left*, average of 6) intensity of stimulation in control (black), with ZX1 (100μM) and after addition of the NMDAR antagonist D-AP5 (50μM). **C)** Plot of EPSP integral (3^rd^ pulse) in function of stimulus intensity obtained from experiment illustrated in B. Dotted lines represent linear regression to the stimulation intensity values before occurrence non-linear EPSPs. **D)** Recorded EPSPs (6 individual sweeps) for conditions listed in B with constant stimulation intensity. **E)** *Left*: Summary plot of integral of synaptically evoked EPSPs (3^rd^ pulse) at the stimulus intensity defined as the minimum value needed to induce non-linear dendritic behavior (in zinc chelated) in control conditions and in the presence of ZX1. *p<0.05. *Right*: the first peak amplitude is not affected by the application of the zinc chelator (ctrl: 7.82 ± 0.83 mV; ZX1:7.97 ± 0.70 mV; n=11 p=0.68 Wilcoxon matched-pairs signed rank test). **F)** Chelating extracellular zinc induces a shift to the left of the stimulus current–voltage relationship. Stimulation intensity is shown relative to the maximum value used in control for each cell define as 1 ( n =11, * p<0.01 Wilcoxon matched-pairs signed rank test). **G)** 2PLSM image of a basal dendrite from a L2/3 PN with 9 selected glutamate uncaging locations (orange). **H)** Photolysis-evoked EPSPs (pEPSPs) in response to increasing number of laser spot locations in control conditions and in the presence of the zinc chelator ZX1. Right: algebraic sum of individual pEPSPs. **I)** Subthreshold input-output relationship of pEPSPs obtained in control conditions (black) and in the presence of ZX1 (orange) for dendrite illustrated in G-H (upper left). **J)** Summary plot of supralinearity for control conditions and after ZX1 application. **K)** Biophysical model of NMDA zinc modulation at single synapse (see Methods). Zinc binding (*b_Zn_*) is modelled by an increment-and-decay dynamics following glutamatergic events. At a given synaptic event, the current level of zinc binding (*b_Zn_*, black curve) sets the zinc modulation level (m_Zn_, piecewise function in orange) that reduces NMDA conductance according to the factor (1-α*_Zn_·m_Zn_*), where α*_Zn_* accounts for the efficacy of zinc inhibition at full binding (see Methods). **L)** Morphological reconstruction of layer 2/3 PN used for the model(Jiang et al., 2015). The orange dot indicates the location of the synaptic stimulation used to obtain the traces in I. **M-N)** Membrane potential (*top*) and conductance (*bottom*) traces obtained following the activation of an increasing number of recruited synapses (*N_syn_*) for free zinc (black) chelated zinc (dark cyan) and AMPA only (blue) conditions. **O)** *Left*: adding of multiple point of synaptic stimulation over the basal dendrite dendritic tree. *Right*: Half-activation level (N_syn1/2_) and respective 3rd pulse integral measured in zinc free (black) or in zinc chelated (dark cyan) obtained for the point of stimulation reported(*left*).***p<0.0001 Wilcoxon matched-pairs signed rank test.

Overall, our results reveal that vesicular zinc is a powerful pathway and cell-specific modulator of NMDAR function in neocortical microcircuits.

### Vesicular zinc is an endogenous modulator of dendritic integration

Having established that synaptic NMDARs are modulated by activity we tested the relevance of such a mechanism in dendritic computations of L2/3 PNs. For that, we patch-clamped L2/3 PNs in the current-clamp configuration and used 2P-imaging to position a stimulation pipette in the close proximity (5-7 μm) of basal dendrites (Figure 3A). Somatic EPSPs were recorded in response to extracellular stimulation consisting of a train of 3 stimuli at 50Hz (Figure 3B). Focal stimulation was validated using dendritic 2P-calcium (Ca^2+^) imaging (Figure S3, Methods). Progressive increase in the number of activated synapses led to the appearance of a slow component in recorded EPSPs that resulted in a supralinear relationship between the intensity of stimulation and the integral of the last EPSP (Figure 3B-D). Interestingly, interfering with the activity-dependent modulation of synaptic NMDARs through zinc chelation resulted in a shift of the response curve towards low intensities. For the same intensity of stimulation depolarizations were increased in the presence of ZX1 (control: 0.75 ± 0.12 mV/sec; ZX1: 1.21 ± 0.17 mV/sec; n=11, p=0.01 Wilcoxon matched-pairs signed rank test; Figure 3E-F) without noticeable alteration in the amplitude of the first EPSP (Figure 3E). D-AP5 (50 μM) application eliminated the supralinearity, as expected for an NMDAR-dependent process (Branco and Häusser, 2011; Palmer et al., 2014; Polsky et al., 2009); Figure 3B-F).

**Figure 3:**
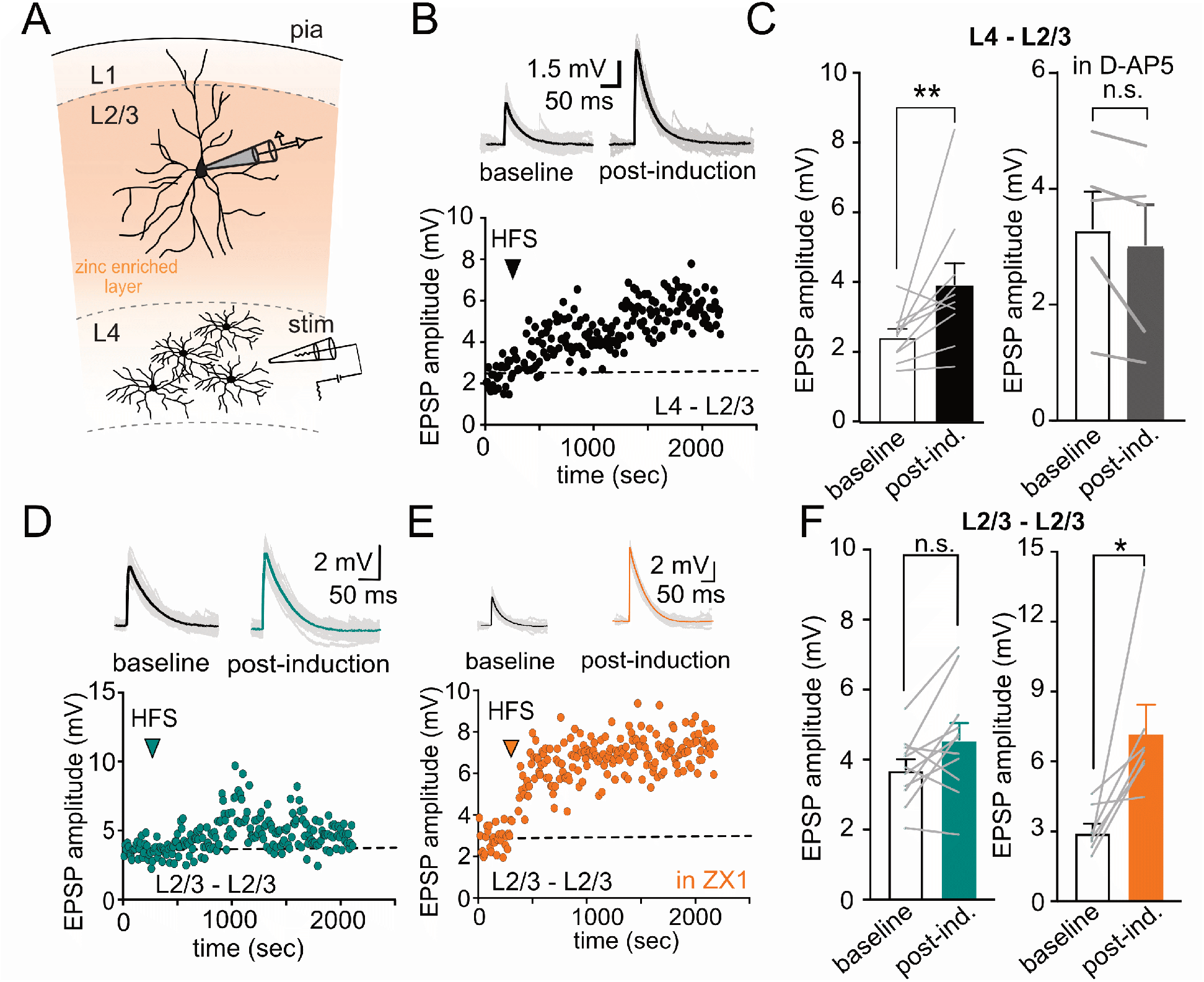
Zinc-mediated reduction of synaptic cooperativity controls LTP induction in a pathway-specific manner in S1. **A)** Schematic representation of experimental conditions used to stimulate L4-L2/3 connections. **B)** Representative traces (up) and amplitude time course (down) of recorded EPSPs in L2/3 PN baseline and after LTP induction while stimulating L4 inputs. **C)** *Left*: Summary plot of EPSPs amplitude, pre- and post-LTP induction in L4-L2/3 connections. *right*: Same as *left* but in presence of D-AP5 (50μM). Data are presented as mean ± SEM. (Layer 4 to 2/3, baseline: 2.44 ± 0.22 mV, post. ind.: 3.94 ± 0.60 mV, p=0.001; Layer 4 to 2/3 D-AP5, baseline: 3.31 ± 0.65 mV, post. ind.:3.03 ± 0.71mV p=0.12; ** p < 0.01, Wilcoxon matched-pairs signed rank test). **D)** Same that B but with LTP protocol applied locally in L2/3-L2/3 connections. **E)** Same as D but for a cell recorded in the presence of ZX1. **F)** Summary plot of average EPSPs amplitudes for total experiments listed in D and E (Layer 2/3 to 2/3, baseline: 3.72 ± 0.28 mV, post ind.: 4.57 ± 0.47 mV p = 0.07; Layer 2/3 to 2/3 ZX1, baseline: 2.97 ± 0.37 mV, post ind.:7.24 ± 1.2 mV p = 0.01; Wilcoxon matched-pairs signed rank test).

The observed modulation of NMDARs by synaptic stimulation would predict no effect of ZX1 in dendritic function if synaptic release is not engaged. To test such prediction, we studied dendritic integration properties of L2/3 PN using 2P-glutamate uncaging. We observed that the EPSP peak amplitude increased with the number of activated synapses closely following a sigmoid function in agreement with the non-linear integration properties of L2/3 PN dendrites (Figure 3G-I; Branco and Häusser, 2011). Under these experimental conditions application of ZX1 did not alter the non-linear behavior of the dendrites tested (Figure 3J, control: 22.8 ± 4.4%; ZX1: 21.8 ± 3.8%, n = 6; p = 0.56 Wilcoxon matched-pairs signed rank test) nor the threshold estimated as the number of inputs required to reach half of the maximum EPSP amplitude (ctrl: 4.33 ± 0.49 inputs; ZX1: 4.50 ± 0.56 inputs; n=6, p > 0.99 Wilcoxon matched-pairs signed rank test). These results argue that during synaptic activation the engagement of endogenous modulatory mechanisms selective for NMDARs function can alter dendritic integration properties of L2/3 PNs.

To further test if modulation of NMDARs could be indeed at the origin of the observed alteration in dendritic activity during ZX1 application we derived a theoretical description of zinc activity-dependent action on NMDARs (Figure 2K, Methods) that we incorporated into numerical simulations of a morphologically-detailed L2/3 PN model (Figure 2L, Methods). Parameters were constrained on experimental measurements (see Methods, Table 1 and Figure S3D-E). To study how modulation of NMDARs by zinc affected dendritic integration, we stimulated synapses with a train of 3 pulses to mimic the experiment of Figure 2B-D and progressively increased the number of activated synapses. In agreement with previous reports and with our own experimental data, model simulation revealed that increased numbers of activated synapses results in a nonlinear NMDAR-dependent increase in measured somatic voltage (Figure 2L-N). Incorporating zinc modulation of NMDARs induced a shift in the input-output curve towards higher stimulus levels (number of synapses needed to reach half-activation voltage, N_syn_^1/2^ = 5.9 ± 1.0 control versus N_syn_^1/2^ in zinc free = 7.0 ± 1.2, p=5e-8, n=25 locations, Wilcoxon matched-pair signed rank test; Figure 2L-O). In parallel, we observed a decrease in the somatically measured depolarization (V_m_) for a given level of synaptic recruitment (3^rd^ pulse integral at N_syn_^1/2^ of the “chelated” condition was 1.4 ± 0.3 mV.s for the chelated zinc condition versus 0.9 ± 0.2 mV.s for the free zinc condition, p=5e-8, n=25 locations, Wilcoxon matched-pair signed rank test; Figure 2M-O).

Altogether, experimental and numerical analysis highlight that endogenous modulation of synaptic NMDARs during repeated synaptic activation regulates dendritic integrative properties of single neurons.

### Activity-dependent modulation of NMDARs and relevance for synaptic plasticity

Activation of NMDARs as long-been associated with plasticity of glutamatergic inputs in neuronal networks. In addition, in L2/3 PNs the recruitment of NMDAR-dependent dendritic nonlinearities have been shown to play an important role in synaptic plasticity both *in vivo* and *in vitro*(Gambino et al., 2014; Williams and Holtmaat, 2019). We thus hypothesized that the segregation of zinc modulation of NMDARs in cortical microcircuits would result in heterogeneity in dendritic integration with the consequent modification of plasticity rules among L2/3 PNs inputs. To test such hypothesis, we applied a high-frequency stimulation protocol, adapted from *in vivo* recordings (see Methods), to either L4 or L2/3 inputs and recorded the subsequent effect in amplitude of evoked EPSPs in L2/3 PNs. As previously reported, LTP was clearly observed at L4 inputs (Figure 3A-C; Williams and Holtmaat, 2019), an effect that was abolished in the presence of the NMDAR antagonist D-AP5. In contrast, the same protocol was ineffective in inducing LTP when local L2/3 axons were stimulated (Figure 3D-F).

In order to test if removing endogenous modulation of NMDARs by zinc could render L2/3 synapses sensitive to such LTP protocols we repeated experiments in the presence of the zinc chelator ZX1 or in ZnT3 KO slices. Interestingly a clear NMDAR-dependent LTP could now be observed in L2/3 pathway (Figure 3E-F, Figure S4A-D). The present results thus reveal that the segregation of vesicular zinc across cortical inputs results in differential recruitment of NMDARs during repetitive synaptic stimulation generating heterogeneity in plasticity induction rules between inputs.

### Zinc modulation of NMDAR implements a background-invariant encoding of synaptic activation under in vivo-like activity

Finally, we investigated theoretically the physiological relevance of the uncovered pathway-specific activity-dependent modulation of NMDARs by zinc in terms of synaptic processing in cortical circuits. Specifically, we analyzed how zinc modulation affects the coincidence detection properties of single dendritic branches under different levels of background activity as observed in vivo (Figure 4). We first calibrated zinc modulation in the model on our experimental data obtained from unitary connections (see Methods) and we simulated passive (Figure 4B) and active (Figure 4C, see Methods and Table 1) cellular integration in response to increasing quasi-simultaneous synaptic recruitment (N_syn_) under different background activity levels (v_bg_). Coincidence detection was quantified by the integrated amount of V_m_ depolarization (PSP integral, Figure 4D) and spike probability (Figure 4E) following the synaptic activation pattern. Both V_m_ and spiking output were strongly influenced by NMDAR activation (Figure 4; note the difference with the “AMPA only” setting). When NMDAR function was simulated without zinc modulation as found in the L4-2/3 connections, coincidence detection was facilitated by increasing levels of background activity (Farinella et al., 2014). Both the V_m_ response and the spiking output exhibited a strong shift toward a lower number of synapses required to reach a similar depolarization/spiking value (Figure 4D and Figure 4E).

**Figure 4.**
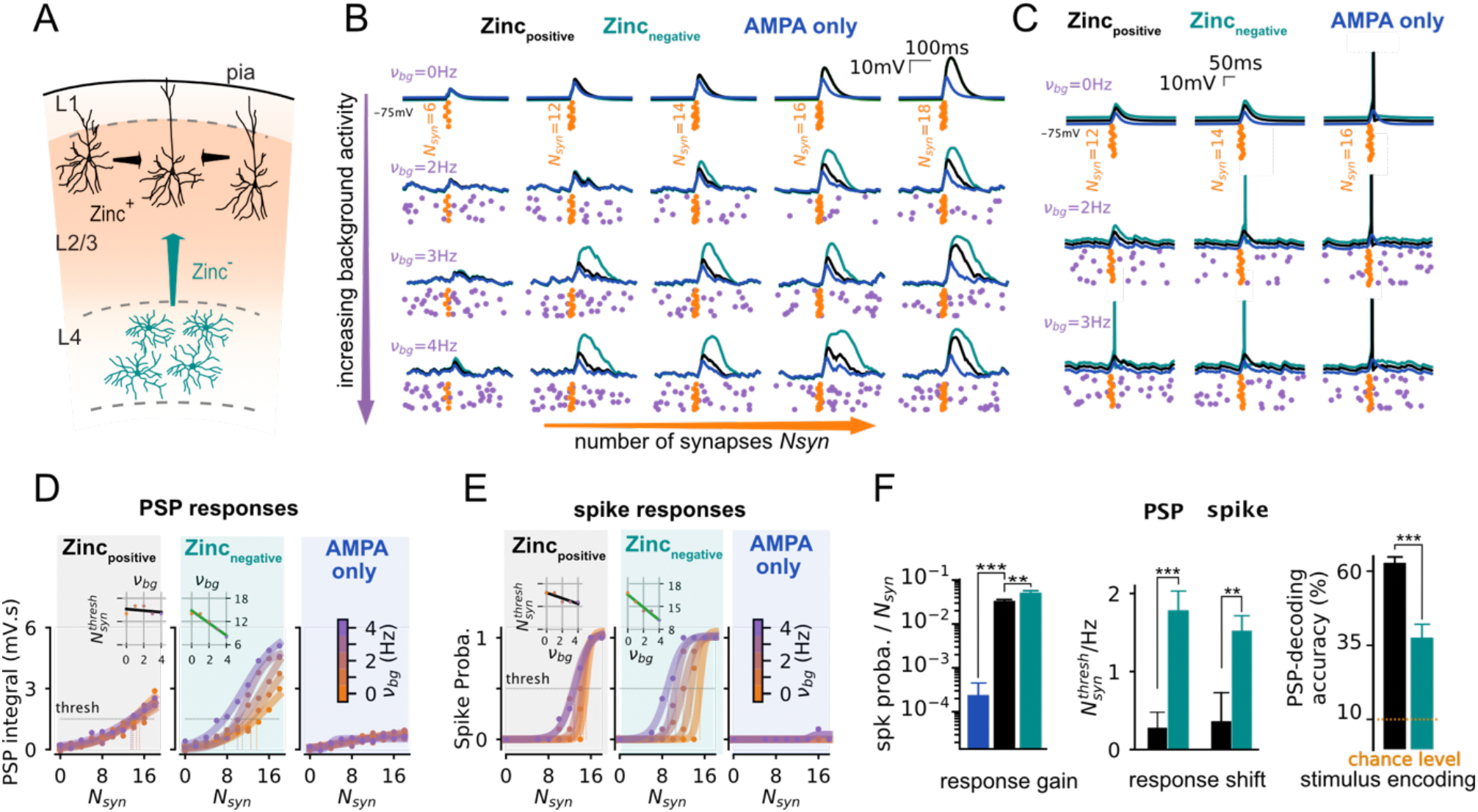
Activity dependent-modulation of NMDARs by vesicular zinc confers unique integrative properties to the L2/3 intralaminar pathway: background-invariant coincidence detection under *in vivo*-like regimes. **(A)** Schematic illustration of zinc-positive (zinc^+^) and zinc-negative (zinc^-^) input pathways into L2/3 PNs. L2/3-to-L2/3 connections have strong zinc modulation (black) while L4-to-L2/3 synapses display no zinc modulation (green). Color code of zinc actions is maintained through figure. **(B)** Examples of single trial somatic *V_m_* traces for subthreshold integration (i.e. simulating passive + synaptic properties only, see Methods) of different numbers of coincidently active synapses *N_syn_* (orange, increasing from left to right) at different levels of ongoing synaptic activity *v_bg_* (increasing from top to bottom). **(C)** Same than **B** but with active conductances added to the model (see **Methods**). **(D)** Integral of PSP responses as a function of synaptic stimulation (*N_syn_*) for the different levels of ongoing synaptic activity (color-coded) at a given synaptic location (response averaged over n=10 with different background activity realisation and n=3 patterns of synaptic stimulations). Data points (dots) were fitted with a sigmoid curve (plain lines). In the L2/3-to-L2/3 and L4-to-L2/3 cases, we highlight the shift of the *N_syn_* level *N_synt_^thresh^* corresponding to a threshold level “*thresh*” (*thresh*=1.5 mV.s) with dashed lines (color-code matching *v_bg_* level). **(E)** Spiking probability response as a function of synaptic stimulation (*N_syn_*) for the different levels of ongoing synaptic activity (color-coded) at a given synaptic location (the spiking probability is computed by averaging over n=10 with different background activity realisation and n=3 patterns of synaptic stimulations). Data points (dots) were fitted with a sigmoid curve (plain lines). We highlight the shift of the *N_syn_* level *N_synt_^thresh^* corresponding to the threshold level “*thresh*” (*thresh*=0.5) with dashed lines (color-code matching *v_bg_* level). **(F)** *Left*: Average gain over background levels (*v_bg_*, the n=5 increasing levels shown in **D**) of the spiking responses for the AMPA-only (blue), L2/3-to-L2/3 (grey) and L4-to-L2/3 (green) synaptic properties, showing mean ± s.e.m. over n=10 synaptic locations. *Middle*: Average responses gain over background levels (*v_bg_*, the n=5 increasing levels shown in **D**) of the spiking responses for the AMPA-only (blue), L2/3-to-L2/3 (grey) and L4-to-L2/3 (green) synaptic properties, showing mean ± s.e.m. over n=10 synaptic locations. *Right*: Performance of a nearest-neighbor decoder (see **Methods**) inferring the synaptic stimulation level *N_syn_* from single trial PSP waveforms for the L2/3-to-L2/3 (grey) and L4-to-L2/3 (green) synaptic properties. *N_syn_* is varied from 0 to 18 in steps of 2 synapses (i.e. 10 steps, see **D,E**) resulting in a chance level of 10% (red bottom line). Showing mean±s.e.m over n=10 synaptic locations and n=3 stimulation patterns (n=30).

The dependency of the threshold crossing level N_syn_^thres^ (see methods) was indeed significantly modulated by the background activity level (Figure 4F) both for the V_m_ response (N_syn_^thres^/ v_bg_=1.8 ± 0.3 Hz^-1^, p = 0.002; Wilcoxon matched-pair signed rank test; Figure 4D) and the spiking output (N_syn_^thres^/ v_bg_ =1.5 ± 0.2 Hz-^1^, p = 0.002, Wilcoxon test; Figure 4E). Strikingly, this facilitation effect disappeared when simulating a zinc containing pathway like the L2/3-L2/3 connections.

Both the V_m_ response and the spiking output lost their strong dependency to the background activity level (V_m_: N_syn_^thres^/ v_bg_ =0.3 ± 0.2 Hz^-1^; p = 2e-3; Figure 4D, F; Spiking: N_syn_^thres^/ v_bg_ = 0.3 ± 0.3 Hz^-1^, p = 0.002, two-sampled Wilcoxon test with zinc-free condition; Figure 4E, F). We therefore addressed the impact in terms of encoding properties of such a phenomenon. We built a decoder of stimulus intensity from the single trial PSP waveform (see Methods). This analysis revealed that the background-facilitation had a confounding effect in the zinc-independent pathway, where a given PSP response could be attributed to different N_syn_ level due the background modulation of evoked activity (*e.g*. a PSP at N_syn_ = 14 and v_bg_ = 3Hz can be confounded with a PSP at N_syn_ = 12 and v_bg_ = 3Hz, see Figure 4F), leading to an overall decoding accuracy of 37.48 ± 4.86 %. This confounding effect disappeared in the putative L2/3-to-L2/3 pathway, the zinc modulation maintained the input-output relationship invariant across background levels what led to a strong and significantly increased decoding accuracy (62.71 ± 2.40 %, p=1.7e-6, Wilcoxon matched-pair signed rank test). We conclude that vesicular zinc release confers unique integrative properties to L2/3-L2/3 synapses, they benefit from the high NMDAR-mediated sensitivity allowing coincidence detection for a few synaptic inputs together with the zinc-mediated NMDAR inhibition that preserves stimulus sensitivity across background activity levels (Figure 4D,E,F).

## Discussion

Synaptic inputs into single cortical neurons exhibit substantial heterogeneity in both structural and functional features. Such a diversity is thought to contribute to the brain’s remarkable information processing capabilities through mechanisms that are still under investigation. We now report that the heterogeneous distribution of vesicular zinc across cortical inputs results in pathway- and cell-specific modulation of NMDARs by synaptic activity. The activity-dependent inhibition of NMDARs shapes dendritic integration properties of cortical neurons and gates synaptic plasticity in an input selective manner. Moreover, experimentally constrained numerical simulations highlighted a previously unnoticed role of NMDARs plasticity in controlling dendritic integration in L2/3 PNs during periods of spontaneous activity like those observed *in vivo*. Inhibition of NMDARs by endogenous zinc normalized dendritic integration by preserving input-output responses for different levels of background activity.

In the neocortex, neurons process information on a varying background of ongoing activity that is associated with the different cortical states (McCormick et al., 2020; Zerlaut et al., 2019). Our numerical simulations suggest that incorporation of spontaneous activity, like observed in awake animals, facilitates dendritic integration of quasi-synchronous inputs (Figure 4). These observations are in agreement with previous reports (Farinella et al., 2014; Ujfalussy et al., 2018) and are largely mediated by increased activation of NMDARs. Facilitated recruitment of NMDA-dependent dendritic non-linearities is particularly evident for distal inputs and is thought to allow closely activated distal synapses to overcome their relative electrotonic disadvantage compared with proximal synapses (Branco and Häusser, 2011). Yet, inputs *in vivo* display heterogenous spontaneous firing rates and the facilitated recruitment of NMDARs is expected to result in a decreased ability of cortical neurons to extract relevant information from inputs displaying lower spontaneous activity (Abbott, 1997). The now reported activity-dependent modulation of NMDARs by vesicular zinc counteracts such effects and contributes to a normalization of synaptic NMDAR participating in dendritic integration across different spontaneous activity regimes. We hypothesize that such a finding might allow neurons to establish a dynamic gain-control mechanism of dendritic non-linearities in face of an heterogeneous background activity.

Computationally, synaptic depression of intracortical synapses has been reported to induce also a frequency-dependent adjustment of synaptic weights allowing a equal percentage rate changes on rapidly and slowly firing afferents to produce equal postsynaptic responses (Abbott et al., 1997). Interestingly, L4-L2/3 PN synapses display pronounced short-term depression and do not have zinc-dependent modulation of NMDARs. In contrast, L2/3-L2/3 PNs synapses have modest synaptic depression (Figure S2G, H) but show clear zinc modulation of NMDARs conductances. Although speculative, these observations suggest that the normalization of synaptic gain in function of afferent firing rates might be a general property of cortical neurons (Carandini and Heeger, 2012) that can be achieved by different cellular mechanisms.

In addition to the direct influence in the input-output transformations performed by single neurons, dendritic operations are also known to control plasticity of synaptic inputs(Gambino et al., 2014; Losonczy et al., 2008). In S1, rhythmic sensory whisker stimulation efficiently induces LTP in L2/3 PNs that depends on the occurrence of NMDAR-dependent dendritic nonlinearities (Gambino et al., 2014). In parallel, both *in vivo* and *in vitro* approaches have observed reduced LTP at horizontal connections across barrel columns in naive mice (Gambino and Holtmaat, 2012; Glazewski et al., 1998; Hardingham et al., 2011). Feed-forward inhibitory mechanisms have been proposed to explain such a difference in plasticity rules (Gambino and Holtmaat, 2012). However, the now observed pathway specific modulation of NMDARs with the consequent increase in threshold for local dendritic non-linearities renders lateral L2/3-L2/3 inputs less sensitive to plasticity protocols when compared with the bottom up L4-L2/3 connections. Such a mechanism is thus expected to reduce LTP at trans-columnar horizontal L2/3-to-L2/3 projections thus contributing to the topographical organization of the barrel cortex by limiting experience dependent modifications of synaptic strength to the barrel associated with the stimulated whisker.

Since the initial discovery of vesicular zinc (Frederickson et al., 2000; Haug, 1967; Maske, 1955) the activity dependent release of this metal ion is thought to shape neuronal activity. However, the impact of zinc containing neurons in the function of neuronal networks remains unclear and poorly understood. An important challenge faced over the years when studying zinc actions in the brain has been the identification of the downstream target(s) of synaptic zinc release. The recent development of a transgenic mouse model carrying a selective impairment in the high-affinity zinc-binding site of the GluN2A NMDA receptor (NMDAR) subunit (Nozaki et al., 2011), revealed that postsynaptic NMDARs are a major target of endogenous vesicular zinc in the hippocampus (Vergnano et al., 2014). Our results in neocortical synapses extend this observation and suggest that at glutamatergic synapses NMDARs are most likely the main target for synaptic zinc release. A striking property of the zinc containing axons is its heterogeneous distribution across the brain(Brown and Dyck, 2004; Frederickson et al., 2000). We now report that such a variability results in input specific alterations in plasticity rules. In addition, our paired recordings between L2/3 PN and local PV-interneurons suggest that zinc modulation of NMDARs can also be adjusted in a postsynaptic manner revealing the modularity of such a mechanism. The underlying cellular mechanism behind the reduced sensitivity of NMDAR in PV-INs is at present unclear but is likely associated with variation in the subunit composition of synaptic NMDARs that can affect the sensitivity of synaptic NMDARs to zinc inhibition (Paoletti, 2011).

Finally, both experimental and numerical analysis provide a new perspective on how zinc shapes neuronal computations. Through an activity-dependent inhibition of NMDARs, vesicular zinc controls dendritic integration in cortical microcircuits and could represent the cellular basis for the altered sensory discrimination observed in ZnT3 KO mice (Patrick Wu and Dyck, 2018).

## Acknowledgements

This study was supported by the Centre National de la Recherche Scientifique, the European Research Council (ERC-STG-678250 to N.R., ERC-ADV-693021 to P.P.), by the Paris Brain Institute, and the Fondation pour la Recherche Médicale (FRM; fellowship ARF 201909009117 to Y. Z. and fellowship FDT201805005817 to B.S.). We thank Dr. Richard Palmiter (University of Washington, USA) for providing the ZnT3-KO mouse line.

## Declaration of Interests

The authors declare no competing interests.

## Supplementary table

**Table 1.** Biophysical and synaptic parameters for the L2/3 pyramidal cell model. Values for the parameters used in the numerical simulations of Fig. 3 and Fig. 4. The parameters highlighted with a star (*) were constrained on experimental recordings. The other parameters were taken from previous studies (Destexhe et al., 1998; Branco and Häusser, 2010; Farinella et al., 2014; Zerlaut and Destexhe, 2017).

| Model parameters | Symbol | Value |  |
| --- | --- | --- | --- |
| Membrane leak conductance* | $G_L$ | 0.29 pS/$\mu\text{m}^2$ | |
| Membrane capacitance* | $C_m$ | 0.91 $\mu\text{F}/\text{cm}^2$ | |
| Intracellular resistance | $R_i$ | 100 $\Omega\cdot\text{cm}$ | |
| Leak reversal potential | $E_L$ | -75 mV | |
| AMPA conductance quantal | $q_{\text{AMPA}}$ | 1.0 nS | |
| AMPA reversal potential | $E_{\text{AMPA}}$ | 0 mV | |
| AMPA rise time | $\tau_{\text{AMPA}}^{\text{rise}}$ | 0.5 ms | |
| AMPA decay time | $\tau_{\text{AMPA}}^{\text{decay}}$ | 5.0 ms | |
| NMDA conductance quantal* | $q_{\text{NMDA}}$ | 2.7 nS | |
| NMDA reversal potential | $E_{\text{NMDA}}$ | 0 mV | |
| NMDA rise time | $\tau_{\text{NMDA}}^{\text{rise}}$ | 3.0 ms | |
| NMDA decay time* | $\tau_{\text{NMDA}}^{\text{decay}}$ | 70.0 ms | |
| Mg<sup>2+</sup> concentration | $[\text{Mg}^{2+}]$ | 1.0 mM | |
| Mg-NMDA sensitivity | $\eta_{\text{Mg}}$ | 0.33 mM<sup>-1</sup> | |
| Mg-NMDA voltage slope factor | $V_{\text{Mg}}^0$ | 12.5 mV | |
| Zinc inhibition efficacy (extracell. stim., Fig. 2)* | $\alpha_{\text{Zn}}$ | 0.19 | |
| Zinc inhibition efficacy (L2/3-L2/3 pairs, Fig. 4)* | $\alpha_{\text{Zn}}$ | 0.45 | |
| Zinc inhibition efficacy (L4-L2/3 extra. stim, Fig. 4)* | $\alpha_{\text{Zn}}$ | 0 | |
| Zinc inhibition decay time* | $\tau_{\text{Zn}}^{\text{decay}}$ | 638.0 ms | |
| **Active current densities per compartment in pS/$\mu\text{m}^2$ (Fig. 4)** |  |  |  |
| **current type** | **basal dendrite** | **soma** | **axon** |
| Sodium channel: “Na”-current | 40 | 1500 | 30000 |
| Potassium channel: “Kv”-current | 30 | 200 | 400 |
| Low-threshold calcium channels | 0 | 0.01 | 0 |
| High-voltage-activation calcium channels: “Ca”-current | 0.5 | 0.5 | 0 |
| Muscarinic potassium channels: “M”-current | 0.05 | 0 | 0 |
| Calcium-dependent potassium channels: “KCa”-current | 2.5 | 2.5 | 0 |

## Supplementary Figures

**Figure S1:**
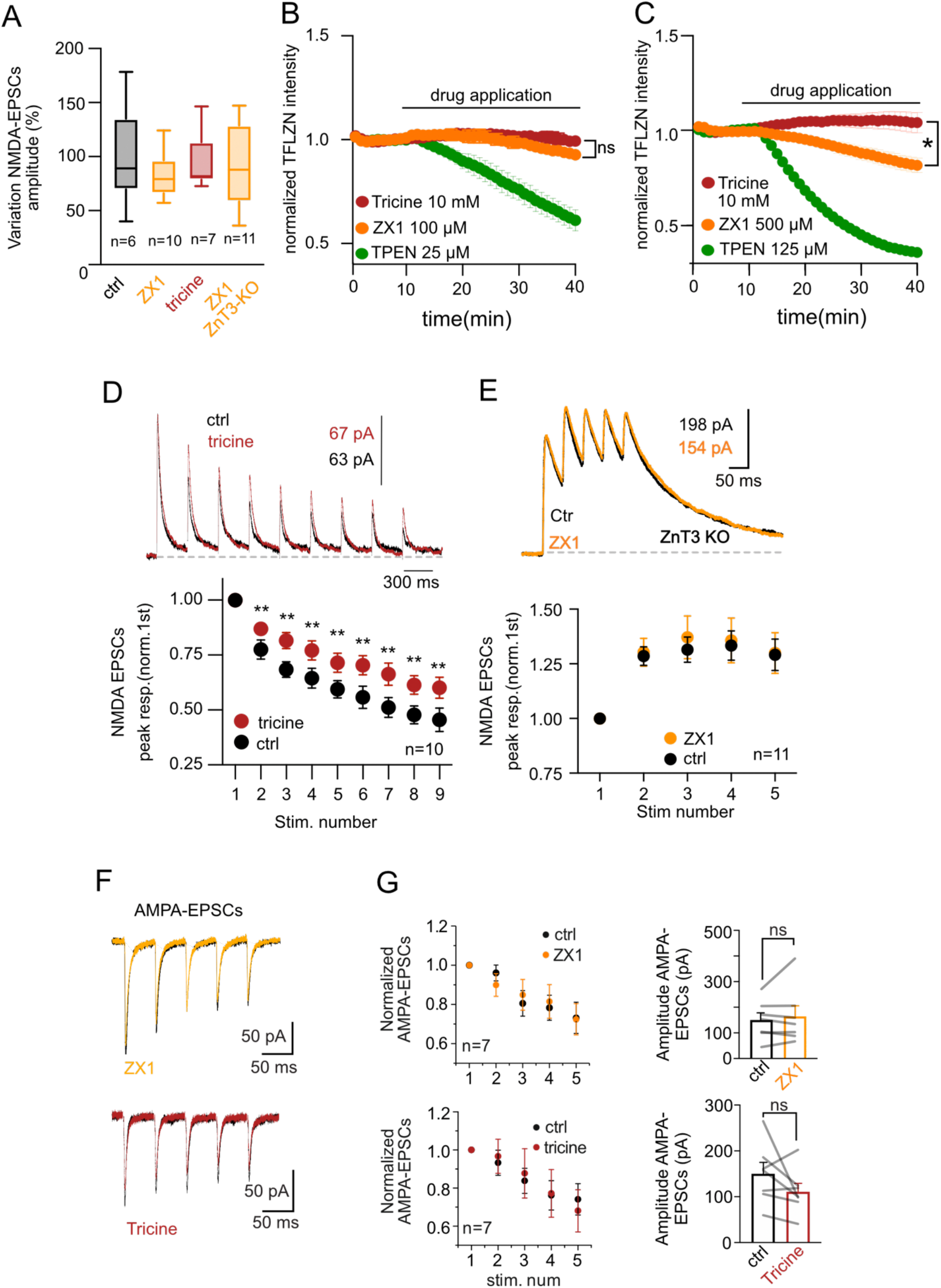
Impact of endogenous zinc in cortical synapses. **A)** Summary plot of percentage change in first peak amplitude of NMDA-EPSCs in control (ctrl, 15 min in ACSF) or during the application of different zinc chelators (15 min application) associated to the experiments in Fig1. ctrl vs chelators P > 0.05 Kruskal-Wallis test. **B)** Normalized TFLZn intensity in stratum lucidum of hippocampus. TPEN application (20 min) but not ZX1 (100 μM) or Tricine (10 mM) significantly reduced TFLZN signal. Tricine n = 4, ZX1 n = 4, PTEN n = 3; p< 0.01 Friedman multiple comparison test. **C)** Same as A but using higher extracellular concentrations of ZX1 (500 μM) and TPEN (125 μM). Note that at higher concentrations, ZX1, and in contrast to Tricine, does reduce intensity of fluorescence of TFLZn suggesting membrane permeation under these working conditions. **D)** *top*: Normalized representative trace of NMDA-EPSCs obtained using train of stimulation consisting of 9 pulses at 3Hz. Bath application of Tricine(10mM) still potentiates NMDA-EPSCs at this frequency of stimulation. *bottom*: Quantification of 10 experiments conducted as illustrated on top. Data are presented as mean SEM (for last pulse, ctrl: 0.45 ± 0.05; Tricine: 0.60 ± 0.05, p = 0.004, Wilcoxon matched-pairs signed rank test). **E)** Potentiation of NMDAR-EPSCs by the zinc chelator ZX1 was absent in brain slices from ZNT3 KO animals, that lack vesicular zinc. Circles indicate mean ± SEM. (for last pulse, ctrl:1.29 ± 0.07; Tricine: 1.30 ± 0.09 p = 0.96, Wilcoxon matched-pairs signed rank test). **F)** Trains of AMPA-EPSCs recorded in control (ctrl) and in the presence of zinc chelators (ZX1 100uM orange, Tricine 10mM red). **G)** Zinc chelation did not affect the amplitude of the first pulse nor the short-term plasticity of AMPA-EPSC trains. p>0.05 Wilcoxon matched-pairs signed rank test.

**Figure S2:**
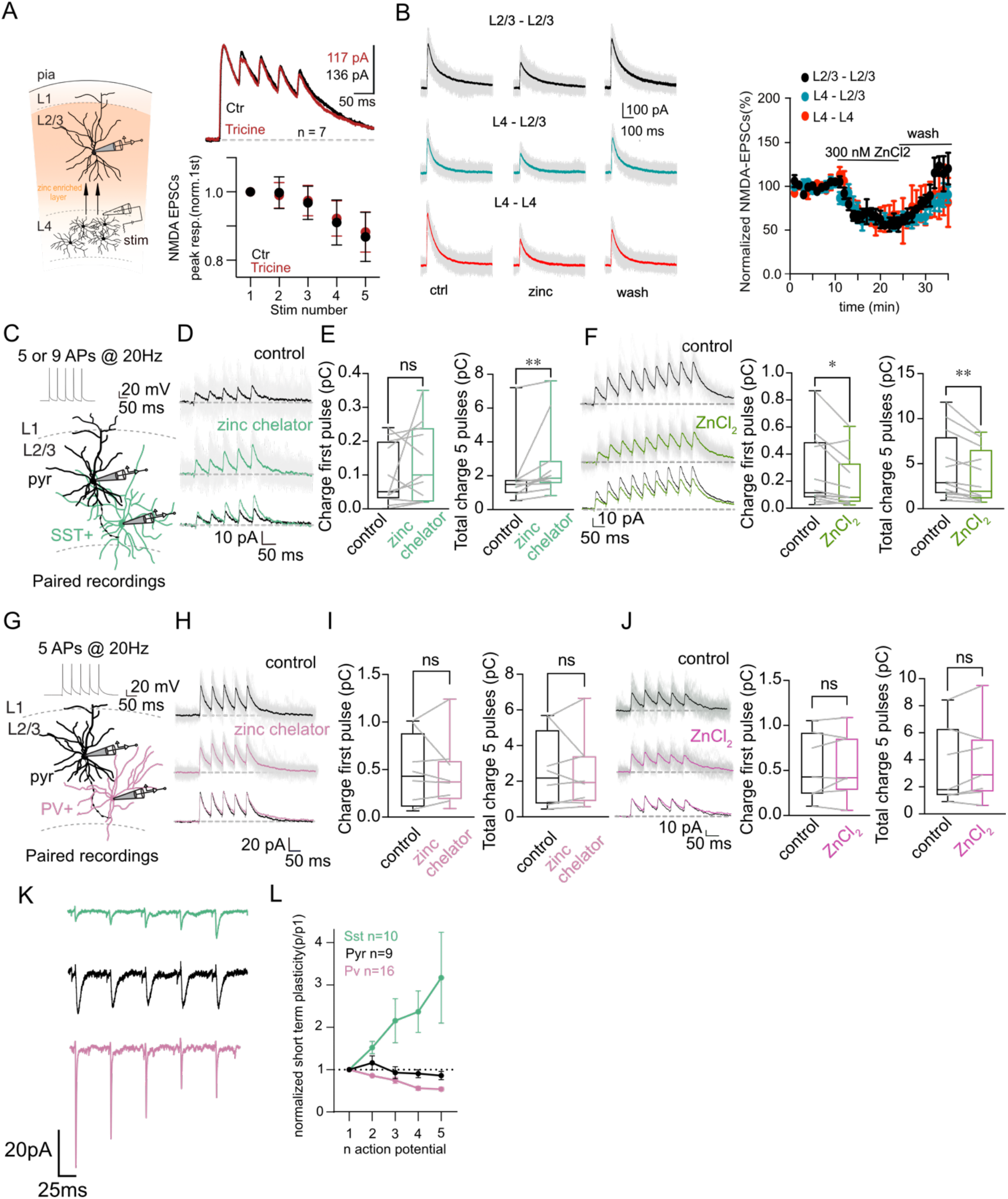
Effect of vesicular zinc in NMDA-EPSCs at L4-L2/3 synapses and at unitary connections within L2/3 neurons of S1. **A)** Schematic representation of experimental conditions used to stimulate axonal inputs from layer 4 and record NMDA-EPSCs in L2/3 PN. Right up: Representative traces of normalized NMDA-EPSCs obtained in control conditions (black line) and after bath application of zinc chelator tricine (red) in L2/3 PNs. Right bottom: Application of the zinc chelator Tricine does not modulate NMDA-EPSCs in the vertical input from L4 to L2/3. Values are presented as mean ± SEM (for last pulse, ctrl: 0.87 ± 0.07, tricine: 0.88 ± 0.06 p = 0.69, paired Wilcoxon matched-pairs test. **B)** *Left*: Effect on exogenous zinc (300nM) application on NMDA-EPSCs while stimulating different excitatory inputs in S1. Color code represents the different synapses studied: L2/3-L2/3 (black), L4-L2/3 (dark cyan), L4-L4 (red). *Right*: Normalized NMDA-EPSCs during zinc application. Max percentage of inhibition: layer 2/3 to 2/3 (n = 4; 56 ± 0.05 %), L4-L2/3 (n = 4; 53 ± 0.04 %), L4-L4 (n = 3; 56 ± 0.12 %). **C)** Illustration of recording configuration used to probe unitary connections between L2/3 PNs and SST^+^-INs. Presynaptic cell was stimulated with a train of 5-9 APs (20 Hz). **D)** Representative traces of uNMDA-EPSCs recorded in a postsynaptic L2/3 SST^+^-IN held at +30 mV. Application of the zinc chelator (green) induced a potentiation of uNMDA-EPSCs. Single repetitions (30 sweeps) are in faint grey, average is in full color. **E)** Summary plot of effect of chelating extracellular zinc in the charge transfer carried by a train of uNMDA-EPSCs. **p<0.001, Wilcoxon matched-pairs signed rank test. Boxes represent interquartile ranges with the horizontal bars showing the medians. **F)** ZnCl_2_ application induced a decrease in the charge transfer of the recorded train of uNMDA-EPSCs in SST^+^-INs. Note that as expected the inhibition by exogenous zinc application is observed already in the first pulse. Data is represented as box plot and quartiles. First pulse charge ctrl: 0.11; ZnCl_2_: 0.08, n=12, p=0.01; total charge ctrl: 2.89, ZnCl_2_: 1.94, n=12, p=0.006, Wilcoxon matched-pairs signed rank test. Representative traces of uNMDA-EPSCs recorded in a L2/3 SST^+^-IN held at +30 mV in ctrl (black) and in the presence of ZnCl_2_ 300 nM (green). **G)** Same as C but for unitary connections between L2/3 PNs and PV^+^-INs. **H-I)** Same as D-E but for uNMDA-EPSCs recorded in a postsynaptic L2/3 PV^+^-IN. **J)** Same as F but for connections between L2/3 PNs and PV^+^-INs. Representative traces of uNMDA-EPSCs recorded in a L2/3 PV^+^-IN held at +30 mV in control (black) and in the presence of ZnCl_2_ 300 nM (dark pink). Data is presented as box plot and quartiles. First peak charge ctrl: 0.43; ZnCl_2_: 0.42 n=7 p=0.58, total charge ctrl: 1.80; ZnCl_2_: 2.89 n=7 p=0.29, Wilcoxon matched-pairs signed rank test. **K)** Example traces of AMPA-uEPSCs obtained in response to a train (5 APs) of presynaptic stimulation of L2/3 PNs (mean 30 sweep) for the different synaptically connected cells in L2/3 of S1. The identity of post-synaptic cell is color coded: SST^+^(green), PN(black) and PV+(pink). **L)** Summary plot of the normalized short-term plasticity obtained for the different post-synaptic targets of L2/3 PNs. Data is presented as mean ± SEM.

**Fig S3:**
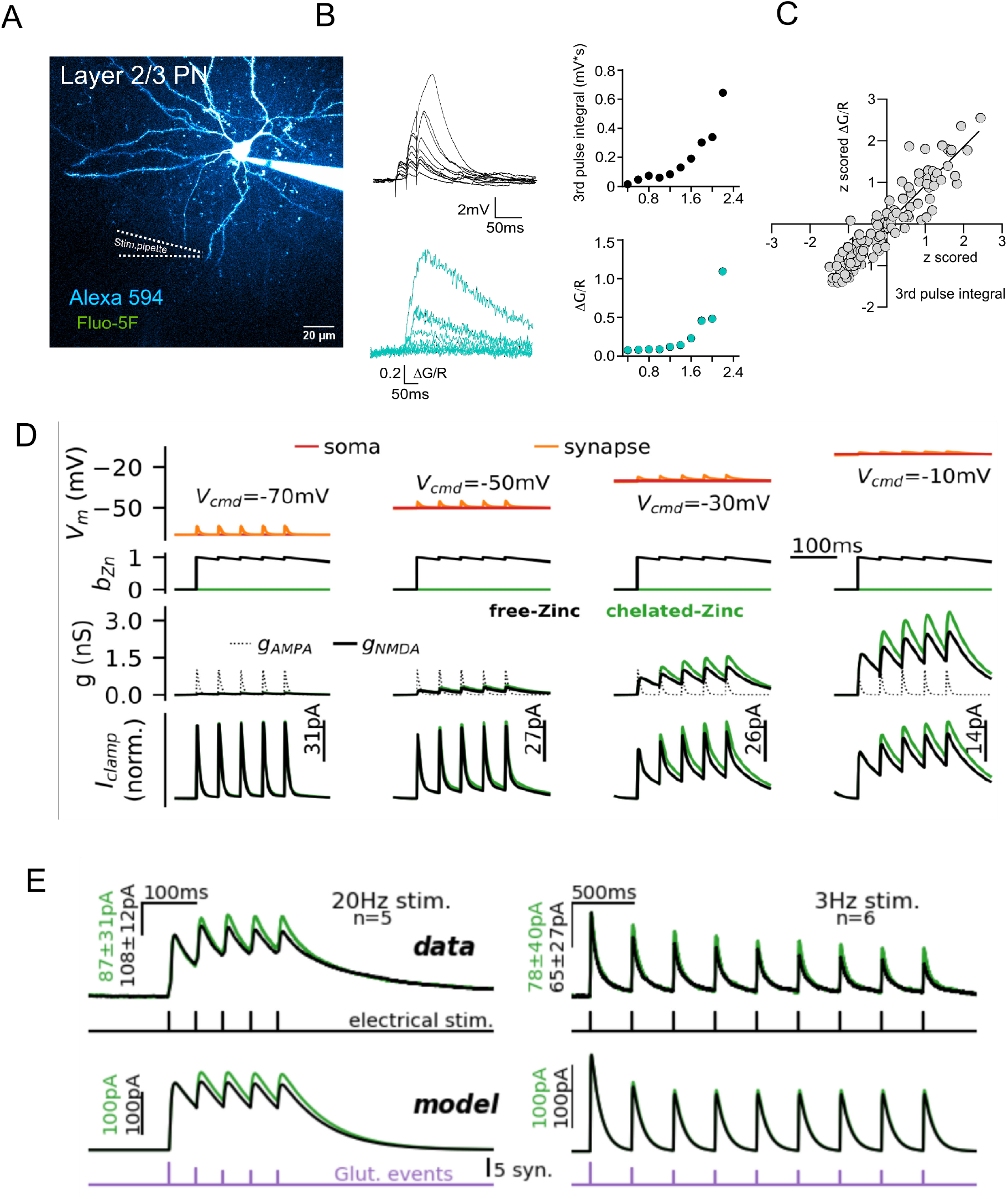
Validation of focal dendritic stimulation and numerical simulation of activity-dependent zinc inhibition of NMDARs. **A)** Two-photon laser scanning microscopy (2PLSM) image (maximum-intensity projection, MIP) of L2/3 PN. **B)** *Top left*: Somatically recorded EPSPs elicited by increasing intensity of stimulation delivered through a glass pipette placed next to dendrite of cell shown in A. *Bottom left*: local dendritic calcium transient associated with EPSPs depicted on top. *Top right*: Plot of EPSP integral (3^rd^ pulse) in function of stimulus current intensity obtained with focal dendritic stimulation. *Bottom right*: Amplitude of calcium transients calculated as ΔG/R in function of stimulus intensity. Amplitude of dendritic calcium transients were highly correlated with the 3^rd^ pulse integral of recorded EPSPs. **C)** Plot of z scored 3^rd^ pulse EPSPs integral and corresponding z scored ΔG/R amplitude obtained from 7 dendrites in n=5 animals. The plot shows a significant correlation p< 0.0001 Person correlation coefficient, R squared= 0.87 between recorded EPSPs and local dendritic calcium signals. **D)** Voltage clamp recording in the model at four different holding potentials following a single synapse stimulation at 20Hz in the presence (black) or absence (chelated) of zinc modulation (green). Reported the membrane potential (top plot: soma in red; dendritic in orange), the evoked conductance at one synapse(middle plot: AMPA and NMDA) and the recorded current at the soma (bottom). **E)** Experimental data used (VC recordings at 20 and 3Hz, top panels) for the optimization of the model parameters. On the bottom panel we show the model response for the optimal set of parameters. Both experimental conditions (free and chelated zinc) were used to optimize the properties of extracellular stimulation in the model and to determine the parameters to simulate zinc modulation (see methods).

**Fig S4:**
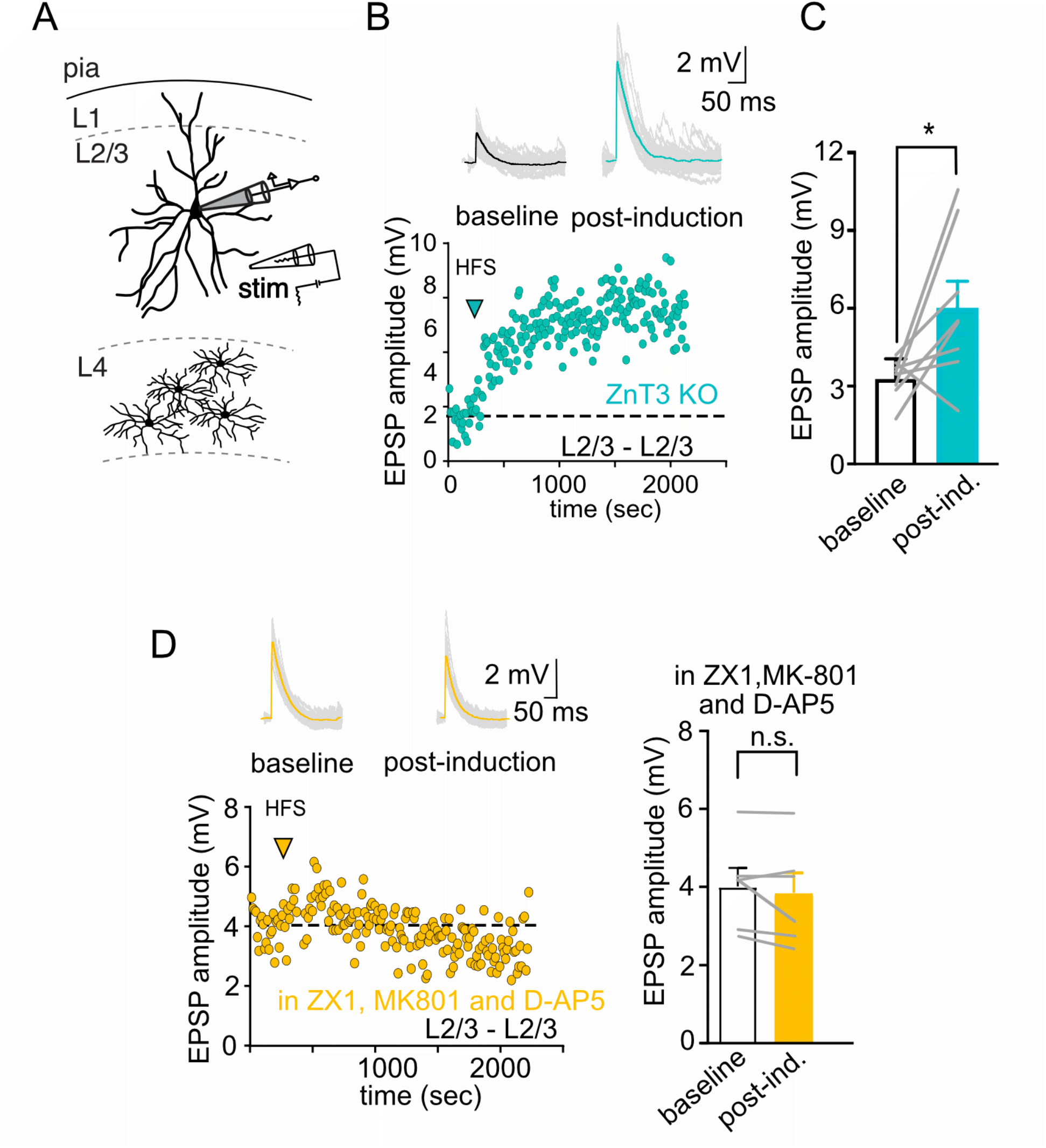
Robust NMDA-dependent LTP in L2/3-L2/3 synaptic connections in brain slices from ZnT3 KO mice. **A)** Schematic representation of experimental conditions used to stimulate L2/3-L2/3 connections. **B)** Example trace and time course of amplitude of recorded EPSPs pre- and post-LTP induction in L2/3 PNs from slices of ZnT3KO mice. **C)** Summary plot of EPSP amplitudes recorded in baseline (pre) and after (post) LTP induction for local L2/3 PNs in ZnT3KO mice. Results are presented as mean ± SEM (baseline: 3.30 ± 0.27, post ind: 6.03 ± 1.02 mV, n=8, p= 0.04, Wilcoxon matched-pairs signed rank test). **D)** Same as B-C, but for experiments performed in the WT mice in the presence of ZX1 and NMDA blockers (MK-801 and D-AP5). Results are presented as mean ± SEM (baseline: 4.03 ± 0.47, post ind:3.87 ± 0.51 mV, n=6, p=0.22, Wilcoxon matched-pairs signed rank test).

## Methods

### Experimental Model

Animals were housed in the Paris Brain Institute animal facility accredited by the French Ministry of Agriculture for performing experiments on live rodents under normal light/dark cycles. Work on animals was performed in compliance with French and European regulations on care and protection of laboratory animals (EC Directive 2010/63, French Law 2013-118, February 6th, 2013). All experiments were approved by local the Ethics Committee #005 and by French Ministry of Research and Innovation. Experimental data was obtained from adult (P21-P41) mice. Both male and female mice were used with the following genotypes: SST-IRES-Cre (SSTtm2.1(cre)Zjh/J; JAX 013044) X Ai9 (Gt(ROSA)26Sortm9(CAG-tdTomato); JAX 007909); PV-Cre ( Pvalbtm1(cre)Arbr/J; JAX 008069) X Ai9 (Gt(ROSA)26Sortm9(CAG-tdTomato); JAX 007909); ZNT3-KO (Palmiter et al.,1996). Animals were maintained on a 12-hour light/dark cycles with food and water provided ad libitum.

### Timm staining

C57BL/6 WT postnatal day (25-35) mice were deeply anesthetized using a ketamine/xylazine mix and transcardially perfused with a perfusate solution prepared by mixing components of the FD Rapid Timm Staining Kit (FD Neurotech). The staining protocol followed the producer protocol with minimum modifications. Modifications: 1)Mice (15-20 gr) were perfused with 50ml of perfusate solution; 2) Brain sections were dehydrated using 50%, 75%, 90%, 100% ethanol for 1 minute each followed by 3 baths of 1 minute in xylene; 3) Sections were then mounted using resinous mounting medium (Eukitt) and glass coverslips.

The slides were scanned using a ZEISS Axio Scan.Z1 device controlled with the ZEISS Software Zen Blue edition. Images were then converted to BigTIFF format and processed with a software programmed in Python 3 using the following libraries: OpenCV, Numpy, Math, Matplotlib, Tkinter. The software detected pixels for which the values match a defined range of colors, thus defining the pixels considered as positive for the staining. Considering the fact that the upper limit of staining has lower RGB values than the lower limit, the use of native pixel values was irrelevant for analysis. Thus the densitometry analysis was achieved by inverting the image’s colors and the boundaries of detection (each RGB values were redefined by subtracting it from the maximum possible value), putting this new image to grayscale and retrieving the gray value of positive pixels (the value of negative pixels was considered as 0).

### Slice preparation

Acute parasagittal slices (320 μm) were prepared from adult C57BL6J mice, starting from postnatal days 25 to 35. For layer 2/3 recordings mice were deeply anesthetized with a mix of ketamine/xylazine (mix of i.p. ketamine [100mg/kg] and xylazine [13 mg/kg]) and perfused transcardially with ice-cold cutting solution containing the following (in mM): 220 Sucrose, 11 Glucose, 2.5 KCl,1.25 NaH_2_PO_4_, 25 NaHCO_3_, 7 MgSO_4_, 0.5 CaCl_2_. After perfusion the brain was quickly removed and slices prepared using a vibratome (Leica VT1200S). Slices containing S1 barrel field were transferred to ACSF solution at 34° C containing (in mM): 125 NaCl, 2.5 KCl, 2 CaCl_2_, 1 MgCl_2_, 1.25 NaH_2_PO_4_, 25 NaHCO_3_, 15 Glucose for 15-20 min. After the period of incubation, slices were kept at room temperature for a period of 6 hours.

### Electrophysiology

Whole-cell patch-clamp recordings were performed close to physiological temperature (33-35°C) using a Multiclamp 700B amplifier (Molecular Devices) and fire-polished thick-walled glass patch electrodes (1.85 mm OD, 0.84 mm ID, World Precision Instruments); 3.5-5 MOhm tip resistance. For voltage clamp recordings, cells were whole-cell patched using following intracellular solution (in mM): 90 Cs-MeSO_3_, 10 EGTA, 40 Hepes, 4 MgCl_2_, 5 QX-314, 2.5 CaCl_2_, 10Na_2_Phosphocreatine, 0.3 MgGTP, 4 Na_2_ATP (300 mOsm pH adjusted to 7.3 using CsOH).

Extracellular synaptic stimulation was achieved by applying voltage pulses (20 μs, 5-50 V; Digitimer Ltd, UK) via a second patch pipette filled with ACSF and placed 20-40 μm from soma. For L4 stimulation, a bipolar concentric stimulation electrode was placed in the “barrel” [visually identified using differential interference contrast (DIC) imaging] below the recorded cell. For current clamp experiments patch pipettes were filled with the following intracellular solution (in mM): 135 K-gluconate, 5 KCl, 10 Hepes, 0.01 EGTA, 10 Na_2_phosphocreatine, 4 MgATP, 0.3 NaGTP (295 mOsm, pH adjusted to 7.3 using KOH). The membrane potential (V_m_) was recorded in current clamp mode (Multiclamp700B amplifier) and held at −70 mV (experimentally estimated RMP = −70.49 ± 1.33 mV, n=23, PNs) if necessary, using small current injection (typically in a range between −50 pA and 200 pA). Recordings were not corrected for liquid junction potential. Series resistance was compensated online by balancing the bridge and compensating pipette capacitance. APs were initiated by brief current injection ranging from 1200-2000 pA and 2-5 ms duration. For calcium imaging experiments Alexa 594 (20 μM,) and the calcium sensitive dye Fluo-5F(300 μM) were added to the intracellular solution daily. For all experiments data were discarded if series resistance, measured with a −5 mV pulse in voltage clamp configuration, was >20 MΩ or changed by more than 20% across the course of an experiment. For current clamp experiment cells were excluded if input resistance varied by more than 25% of the initial value. To record NMDA-EPSCs, picrotoxin (100 μM, Abcam) and NBQX (10 μM, Tocris) were added to the ASCF to block GABAA and AMPA receptors respectively. All recordings were low-pass filtered at 10 kHz and digitized at 100 kHz using an analog-to-digital converter (model NI USB 6259, National Instruments, Austin, TX, USA) and acquired with Nclamp software(Rothman and Silver, 2018) running in Igor PRO (Wavemetrics, Lake Oswego, OR, USA).

### Paired Recordings

For L2/3-L2/3 PN recordings, neurons were identified using DIC imaging. For L2/3-PV-INs and L2/3-SST-INs paired recordings cortical interneurons were identified using PV-Cre and SST-cre mice crossed with the reporter mouse line (Ai9, (ROSA)26Sortm9(CAG-tdTomato)Hze/Jt; JAX 007909). The intracellular solution used for presynaptic cell was (in mM): 130 K-MeSO_3_, 4 MgCl_2_, 10 Hepes, 0.01 EGTA, 4 Na_2_ATP, 0.3 NaGTP (300mOsm, pH 7.3 adjusted with NaOH). For the postsynaptic cell the same solution as described above for voltage clamp was used. Connections were probed using a train of 5/9 action potentials at frequency of 20Hz. After a clear identification of synaptic connection (mean of 30 sweeps with clear AMPA current), post-synaptic cell membrane potential was brought from the initial −70 mV membrane potential to +30mV to record the NMDAR-EPSCs in the presence of blockers as described before. Exogenous zinc application was obtained using a tricine (10mM) buffered ACSF. At pH 7.3 and with 10 mM tricine, calculations show that there is a linear relation, [Zn]_f_ = [Zn]_t_/200 for [Zn]_f_ < 1 μM(Fayyazuddin et al., 2000; Paoletti et al., 1997; Vergnano et al., 2014). A concentration of 60 μM zinc chloride was added to the solutions to obtain 300nM of free zinc.

### LTP protocol

After entering whole-cell configuration a 5 min period of evoked (frequency:0.1 Hz) excitatory post-synaptic potentials (EPSPs) was recorded as baseline. LTP induction protocol consisted of 3 pulses at 100Hz repeated 160 times with and inter stimulus interval of 2.5Hz for a total duration of 1.2 min. This protocol was based on the previous observation that rhythmic whisker stimulation (duration of 1min) induces LTP of glutamatergic L4 inputs into L2/3 cells both *in vitro* and *vivo*. The 3 pulses at 100 Hz were chosen to mimic the train of action potentials (1-3 APs) at high frequency (200-400Hz) observed in L4 neurons in response to whisker stimulation(Yu et al., 2019). Synaptic EPSPs were recorded for an additional 30 min after LTP induction. Cells were discarded if the input resistance varied by more than ± 25% from the initial value and if a ± 5mV change in the resting membrane potential occurred. For the statistical analysis the average of the EPSPs over the baseline period of 5 min (baseline) was compared with mean obtained in the final 5 minutes of the 30min recording after the induction (post-induction).

### TFLZn experiments

For zinc imaging experiments, 21-28 days old C57BL6/J mice (Charles River) were used. After deep anesthesia using isoflurane (Iso-Vet), mice were decapitated and the brain was rapidly removed and transferred in a slicing chamber filled with a cold (4 °C) slicing solution containing sucrose (230 mM), glucose (25 mM), KCl (2.50 mM), NaH_2_PO_4_.H_2_O (1.25 mM), NaHCO_3_ (26 mM), CaCl_2_ (0.8 mM), MgCl_2_ (8 mM) and bubbled with 95% O_2_ and 5% CO_2_. Coronal brain slices 320 μm thick were obtained using a vibratome (Campden Instruments 7000 SMZ2). Slices were then immediately transferred in a chamber containing artificial cerebrospinal fluid (ACSF) at 31-33 °C and recovered 1h before use. ACSF classically contained NaCl (125 mM), KCl (2.5 mM), NaH_2_PO_4_.H_2_O (1.25 mM), NaHCO_3_ (26 mM), CaCl_2_ (2 mM), MgCl2 (1mM) and was bubbled with 95% O_2_ and 5% CO_2_. In order to stain for vesicular zinc, brain slices were incubated for 1h at room temperature in a solution containing ACSF, TFLZn K^+^ salt (250 μM, TEFLabs) and bubbled with 95% O_2_ and 5% CO_2_. Slices were then transferred in the experimental chamber and continuously perfused with a solution at 31-33 °C containing oxygenated ACSF, bicuculline methochloride (10 μM, Hello Bio), tetrodotoxin (0.2 μM, Tocris) and either TPEN (25 μM or 125 μM, Sigma-aldrich), tricine (10 mM, Sigma-aldrich), or ZX1 (100 μM or 500 μM, Strem Chemicals). Excitation was provided by a UV pulse using a 365 nm LED (Thorlabs). UV illumination was concomitant to image acquisition, lasting 700 ms. Frequency acquisition was set at 1 frame per minute. Emitted fluorescence (510 nm) was collected and recorded using an Orca-FLASH4.0 camera (Hamamatsu) mounted on an upright microscope (Scientifica) equipped with a 10x water immersion objective (Olympus). The entire hippocampus was imaged using a motorized stage (Scientifica) control by Micromanager software (v1.4). Images were analyzed using ImageJ software (v 1.52n). Mean fluorescence intensities were measured for each image of a data set after defining a region of interest consisting of a representative area the stratum lucidum.

### 2P Ca^2+^ imaging

Cells were identified and whole-cell patch-clamped using infrared Dodt contrast (Luigs and Neumann, Ratingen, Germany) and a frame transfer CCD camera (Infinity-Lumenera). Two-photon fluorescence imaging was performed with a femtosecond pulsed Ti:Sapphire laser (Cameleon Ultra II, Coherent) tuned to 840 nm coupled into an Ultima laser scanning head (Ultima scanning head, Bruker), mounted on an Olympus BX61WI microscope, and equipped with a water-immersion objective (60X, 1.1 numerical aperture, Olympus Optical, Tokyo, Japan). Cell morphology was visualized using fluorescence imaging of patch-loaded Alexa 594 (20 μM). Ca2+ transients induced by focal synaptic stimulation were recorded using the calcium indicator Fluo-5F (300nM) and rapid line scan imaging (0.76 ms per line). Total laser illumination per sweep lasted 400-600 ms. Fluorescence recordings started at least 25 min after establishing whole cell configuration for layer 2/3 cells, (Fig. S3) to allow dye equilibration. 7-10 linescans were recorded per dendrite (Layer 2/3 PNs) or with an intersweep frequency of 0.33 Hz. Recordings were discarded if red signal decreased/increased by ±20% of the initial value indicating incomplete initial dye equilibration. Fluorescence light was separated from the excitation path through a long pass dichroic (660dcxr; Chroma, USA), split into green and red channels with a second long pass dichroic (575dcxr; Chroma, USA), and cleaned up with band pass filters (hq525/70 and hq607/45; Chroma, USA). Fluorescence was detected using both proximal epi-fluorescence and substage photomultiplier tubes: multi-alkali (R3896, Hamamatsu, Japan) and gallium arsenide phosphide (H7422PA-40 SEL, Hamamatsu) for the red and green channels, respectively.

### Analysis Ca^2+^ imaging

Calcium transients were extracted from linescan images. The fluorescence as a function of time was averaged over visually identified pixels corresponding to width of the dendrite and then averaged over individual trials, resulting in a single fluorescence trace as a function of time (Fdendrite(t)). The background fluorescence (Fback(t)) was estimated similarly (identical spatial line length), but from a location not on a labeled structure and the average value subtracted from Fdendrite(t). Changes in fluorescence were then quantified from the background corrected traces as: ΔG/R(t)=(F_green_(t)-Frest,_green_)/(F_red_) where F_green_(t) is the green fluorescence signal as a function of time, F_rest,green_ is the green fluorescence before stimulation and F_red_ is the average fluorescence of the red indicator (Alexa 594).

### Focal dendritic stimulation

Focal two-photon guided synaptic stimulation in basal dendrites of L2/3 PNs was performed using a theta-glass pipette (series resistance 6-7 MegaOhm). Theta glass pipette filled with extracellular solution was placed in close proximity to the visual identified dendritic segment (5-7micrometer). Experiments were performed in the absence of pharmacological blockers and in the presence of D-Serine (100 μM). Synaptic responses were evoked (20 μs, 0-3 A; Digitimer Ltd, UK) using a short burst of 3 pulses at 50 Hz. The local stimulation was increased (20μAmp per step) until a clear slow component of the EPSP, classically associated with the recruitment of NMDA conductances (NMDA spike) was observed. In a subset of experiments, simultaneous calcium imaging experiment was used to test focal stimulation (see above). Somatically recorded EPSPs were highly correlated with calcium transient imaged in the dendrite close to the stimulation pipette. The threshold for NMDA spike was defined as the stimulating intensity that resulted in the presence of the slow EPSP decay in 50% of trials.

### Glutamate uncaging

L2/3 PN morphology was visualized using fluorescence imaging of patch-loaded Alexa 594 (20 μM). The output of two pulsed Ti:Sapphire (DeepSee, Spectra Physics) lasers were independently modulated to combine uncaging of MNI-glutamate and Alexa 594 imaging. The imaging laser beam was tuned to 840 nm and modulated using a Pockels cell (Conoptics, Danbury, CT). For uncaging, the intensity and duration (500 μs) of illumination of the second Ti:Sapphire laser tuned to 720 nm was modulated using an acousto-optic modulator (AA Opto-Electronic, France). MNI-caged-L-glutamate (20 mM) was dissolved in a solution containing (in mM): NaCl 125, glucose 15, KCl 2.5, HEPES 10, CaCl_2_ 2 and MgCl_2_ 1 (pH 7.3) and constantly puffed applied using a glass pipette (res 2-2.5 MOhm) placed in the proximity of a selected dendrite. Multiple uncaging locations (6-8) were placed adjacent (1 μm) to visually identified spines and not closer than 2 μm to avoid glutamate spillover between locations. Laser intensity was adjusted to obtain individual pEPSP with amplitude between 0.2-0.9 mV (mean = 0.67+/- 0.06 mV). Photolysis laser powers, estimated at the exit of the objective were <20 mW. Interstimulus interval varied between 100-250 ms (isolated pEPSPs) and quasi-simultaneous activation (120 μsec between locations) The arithmetic sum (corrected for not perfect simultaneous spine activation) was used to compare the value obtained (expected) versus the recorded EPSP elicited by a increasing number of spines recruited by glutamate uncaging. An input-output curve was obtained by plotting the amplitude of the expected EPSP versus the amplitude of recorded EPSP. Non-linearity was calculated as previously decribed (Schmidt-Hieber C. et al. 2017):

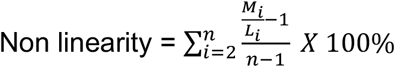

Where n is the maximal number of synapses activated, M_i_ is the amplitude of the measured EPSP, L_i_ is the amplitude of the EPSP constructed by the liner summation of the single synapses taking into account the relative timing of stimulation.

### Computational model of L2/3 pyramidal cell

We performed numerical simulations of single cell integration on morphologically-detailed L2/3 pyramidal cell (ID “L23pyr-j150407a” from the publicly available morphology dataset from Jiang et al., 2015, see **Figure 3F**). Single cell integration was simulated using the cable equation (Koch, 1984; Rall, 1962):

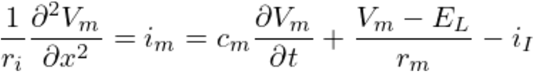

where the membrane current *i_m_* and the input currents *i_I_* are linear density of currents. The cable equation parameters are derived from the membrane parameters of **Table 1** by applying the radial symmetry (for a segment of diameter *D*: *r_m_*=1/(*D·G_L_*), *c_m_*=*π·D·C_m_, r_i_*=(*4·R_i_*)/(*π·D^2^*). Additional point currents (synaptic inputs and/or “voltage-clamp”-currents) are inserted in a segment through the *i_I_* term (as *i_I_*=*I_point_/l_S_* for a current *I_point_* and a segment of length *l_S_*). For voltage-clamp protocols, the leak conductance *g_L_* was decreased by a factor 5 to reproduce the Cesium block, an additional point current *I_clamp_*= *g_clamp_·(V_cmd_-V_m_*) of clamping conductance 1μS was inserted at the soma, an additional 200ms were added prior to stimulation to reach stationary “clamp” conditions (initialized at *V_cmd_*) and we report the *I_clamp_* quantity after those 200ms (see **Figure S3D**). The simulations were implemented using the *Brian2* simulator (Stimberg et al., 2019). Numerical integration was performed with an “exponential Euler” integration scheme and a time step of 0.025ms.

#### Synaptic currents

We considered 2 types of synaptic transmission: NMDA and AMPA. For both synaptic types, the temporal profile of the synaptically-evoked conductance variations was made of a double exponential waveform (Destexhe et al., 1998; Koch, 2004; Zerlaut and Destexhe, 2017).

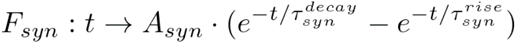

where *τ_syn^rise^_* and *T_syn^decay^_* are the rise and a decay time constant respectively. The waveform was normalized to peak level with the factor:

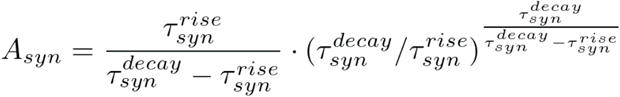

For an AMPA synapse activated by a set of events *{t^k^}* located at position *i* (of membrane potential *V_m^i^_*), the synaptic current *I_syn^i^_* reads:

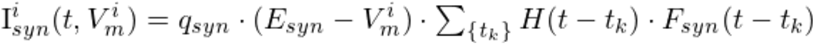

where *q_syn_* is the conductance quantal of the synaptic release (setting the peak conductance level) and *H(t)* is the Heaviside (step) function. We relied on our experimental measurements to determine the ratio of NMDA to AMPA conductances. We found the ratio of NMDA to AMPA peak currents to be 2.7±1.5 (n=10 cells), similar to previous reports (Branco and Häusser, 2011). This ratio was used to set the NMDA conductance quantal from the AMPA conductance quantal (see **Table 1**).

For NMDA synapses, we add a voltage-dependency due to the Magnesium block captured by an exponential function (reviewed in Koch, 2004) and a zinc-binding dependency (detailed in the next section):

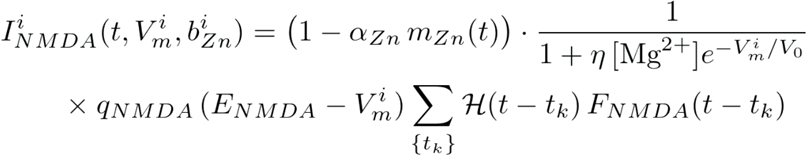

The synaptic parameters are summarized on **Table 1**.

#### Modeling zinc-modulation of NMDAR signaling

We derive a model of the zinc modulation of NMDAR signaling based on our experimental observations and previous studies (Vergnano et al., 2014). Our modeling approach aimed at describing the modulation in the form of a minimal phenomenological description at the single synapse level. The reasoning guiding the derivation of the model is the following (illustrated in **Figure 2K**). Glutamatergic events are associated to a release of zinc in the synaptic cleft. zinc has a high affinity for binding and does not accumulate in the cleft (the cleft is free of zinc in less than 2ms), so the dynamics is led by the binding-unbinding phenomenon. zinc binding is modelled by a variable *b_Zn_* varying betweeen 0 (no binding) and 1 (full binding). zinc binding is approximated as instantaneous and fully saturating the binding site at every single synaptic release event, i.e. it is updated to fully-bound level *b_Zn_*,=1 at each glutamatergic event. zinc unbinding has a decay time constant T_Zn^decay^_ The dynamics of zinc binding *b_Zn_* therefore follows the equation:

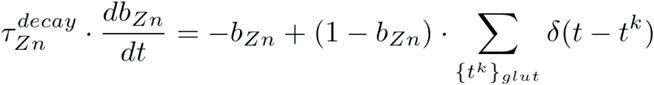

At the time of a synaptic release, the current level of zinc binding will determine the magnitude of the reduction of NMDA conductance due to the zinc modulation. The factor modulating the NMDA conductance *m_zn_(t)* is a piecewise time-varying function updated to the *b_zn_* level at every synaptic release. Therefore, for a set of glutamatergic events *{t^k^}*, the time-varying factor is defined by the piecewise function:

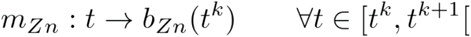

Finally, a parameter *α_Zn_* captures the inhibitory efficacy of zinc binding on NMDAR conductance. At full binding (*b_zn_*=1), the NMDA conductance has a factor 1-*α_zn_*. The zinc modulation is therefore summarized by the factor 1 - *α_Zn_·m_zn_(t)* (see previous section). The dynamics of the *b_Zn_* and *m_Zn_* variables following synaptic events and their impact on the time-varying conductance *g_NMDA_(t)* is illustrated on **Figure 2K** and we describe the model behavior over different *V_m_* ranges in **Figure 2K**.

#### Synaptic locations on the basal dendrites

To host the synaptic stimulation, we looked for n=25 locations (set of segments) distributed over the basal dendrite. The criteria to include a location was that the set of segment should be contiguous over the dendritic tree (i.e. not spread over multiple branches) and the starting point should at least 50μm far from the soma (as excitatory synapses tend to avoid perisomatic locations). We looped over random starting segments in the dendritic tree, a location was considered as the starting segment with the next 20 following segments. The location was included if the criteria was matched and the loop was stopped when we found n=25 locations. The resulting set of the synaptic locations is shown in the inset of **Figure 2O**. We then spread one synapse per segment. From this set of locations, we can therefore recruit up to 20 synapses (see **Figure 2O** or **Figure 4**).

#### Model calibration: parameter fitting

The parameter of the model were fitted in successive steps.

##### Passive properties

To calibrate the passive properties of the model, i.e. the membrane leak conductance *G_L_* and the membrane capacitance *C_m_* (see **Table 1**), we used our set of current-clamp experimental recordings to determine the input resistance and capacitance at the soma. The experimental values were found to be *R_soma_*=137.5±50.2 MΩ, *C_soma_*= 208.1±43.0 pF (n=27 cells). In the model, we varied the passive parameters and computed the input resistance and capacitance at the soma. This was achieved by fitting the Vm response to the single compartment response following a 200 pA current step. The range for the parameter parameter variation was *G_L_* in [0.02,2.0] pS/μm^2^ and *C_m_* in [0.5,2.0] μF/cm^2^ on a [30×30] regular grid. The joint minimization (via the product of the normalized square residual) of the somatic *R_soma_* and *C_soma_* in the model led to the parameters *G_L_*=0.29 pS/μm2 and *C_m_*=0.91 μF/cm2, similar to previous analysis of layer 2/3 pyramidal cell in mice sensory cortex (Branco and Häusser, 2011; Palmer et al., 2014).

##### Biophysical parameters of zinc modulation

We used our set of voltage-clamp recordings with extracellular stimulation at 20Hz and 3Hz in L2/3 (data of **Figure 1B,C**) to constraint the zinc-dependent NMDAR model. We optimized the product of the normalized residual traces at 20Hz and 3Hz between model and experiment. The experimental data was a “grand-average” over cells after trial-averaging (see “data” in **Figure S3E**). The first step consisted in using the “chelated-zinc” condition to evaluate the NMDAR kinetics (*T^decay^_NMDA_*) and to find the correlate in the model of the extracellular stimulation protocol. We described the electrical stimulation as a number of activated synapses *N_syn_* decreasing with stimulation time (to capture the decreasing efficacy of the electrical stimulation in the control case, see **Figure S3E**): *N_syn_(t)*= (*N_syn^0^_*-*N_syn^∞^_*)·*exp*(-*t/τ_N_*)+N_syn^∞^_. We varied the parameters in the range *T^decay^_NMDA_* in [60, 120]ms, *N_syn^0^_* in [3, 11] synapses, *N_syn^∞^_* in [2, 10] synapses and τ_N_ in [30,70]ms on a [7×5×5×4] linear grid. The best fit model was found for: *T^decay^_NMDA_*=70.0 ms, *N_syn^0^_*=7 synapses, *N_syn^∞^_*=2 synapses and τ_N_=56.7 ms (see “chelated-zinc” in **Figure S3E**). In a second step, we used the “free-zinc condition” to determine the zinc-modulation parameters of the NMDAR model introduced (*α_zn_* and *T^decay^_Zn_*). We varied the zinc parameters in the range *α_Zn_* in [0.1, 0.6] and *τ^decay^_Zn_* in [100,1300] ms on a [30×30] linear grid. The best fit model was found for: *α_Zn_* =0.19 and *τ^decay^_Zn_* =638.0 ms.

##### Pathway specificity of zinc modulation

To estimate the parameters of zinc modulation corresponding to each synaptic pathway, we first computed the relationship in the model between zinc efficacy *α_Zn_* and the level of charge increase under zinc chelation (i.e. with respect to the *α_Zn_*=0 setting) in the fifth pulse following a 5 pulse stimulation at 20Hz (see **Figure 1** and **Figure S3D**). We then inverted this relationship to relate an experimental observation to a model setting. For the L4-to-L2/3 pathway, the zinc chelation effect following extracellular stimulation in L4 was non-significantly deviating from 0 (**Figure S2A**) so the corresponding zinc efficacy was set to *α_Zn_*=0. For the L23-to-L23 pathway, zinc chelation increased charged by 47±16% in paired intracellular recordings (**Figure 1E, F**) and the corresponding level of zinc efficacy in the model was found to be *α_Zn_*=0.45.

#### Model of background and stimulus-evoked synaptic activity

Background synaptic activity was adapted from a previous study (Zerlaut and Destexhe, 2017). At each synapse, a given rate value v_bg_ was converted into a presynaptic pattern thanks to an homogeneous Poisson point process. All simulations were repeated over n=10 different realisations of the background activity. Stimulus-evoked activity was designed as follows. For a stimulation at time *t* with a level of synaptic recruitement *N_syn_*, we randomly chose *N_syn_* synapses over the 20 synapses available at each synaptic location (see above) and the event of each of the *N_syn_*, synapse was drawn from a uniform distribution of 20 ms width centered at *t*. The simulations were repeated over different realisations of the stimulus pattern. Example of the background and stimulus-evoked synaptic events for single trials are shown on **Figure 4B,C**.

#### Active membrane currents

We analyzed the impact of active cellular mechanisms (**Figure 4**) by adding the following currents in the single cell model: “Na”-current, “Kv”-current, “T”-current, “Ca”-current, “M”-current, “H”-current, “KCa”-current and a decay dynamics for the Calcium concentration (reviewed in Koch, 2004). The currents and their densities in the different compartments are listed **Table 1**.

### Decoding synaptic stimulation level from single trial *V_m_* responses

We built a decoder of the stimulus intensity classifying single-trial somatic V_m_ responses in the [−100,300] ms interval surrounding stimulus onset (**Figure 4**). The waveforms were baseline-subtracted (estimated from the [−100,0] ms interval) to remove the influence of the relationship between background activity and baseline level. The training set was defined as the set of the trial-averaged response (averaging over n=10 background realisations for each synaptic location and stimulus seed) and the test set was the set of single trial responses for a given synaptic location and stimulus seed. The decoder was implemented using the *NearestNeighbors* function of *sklearn* (Pedregosa et al., 2011) and we used the *euclidean* metric to evaluate distance between response waveforms.

### Statistical analysis

Data analysis was performed using IGOR Pro (Wave Metrics).All the statistical analysis was performed and graphs were created using Prism 8 (GraphPad). Most data are represented as mean ± SEM or using box and scatterplots depicting the median, the interquartile ranges as well as individual values. For statistical analysis normal distribution was not assumed and the following non-parametric tests were used: Wilcoxon matched-pair signed rank test for paired samples, Mann-Whitney test for unpaired, Kruskal Wallis test followed by Dunn’s test to correct for multiple comparison if necessary. Significance was conventionally set as ***p < 0.001, **p < 0.01 and *p < 0.05.

## Notes

### Competing Interest Statement

The authors have declared no competing interest.

